# Dynamic modulation of enhancer responsiveness by core promoter elements in living *Drosophila* embryo

**DOI:** 10.1101/2021.03.18.435761

**Authors:** Moe Yokoshi, Manuel Cambón, Takashi Fukaya

## Abstract

Regulatory interactions between enhancers and core promoters are fundamental for the temporal and spatial specificity of gene expression in development. The central role of core promoters is to initiate productive transcription in response to enhancer’s activation cues. However, it has not been systematically assessed how individual core promoter elements affect the induction of transcriptional bursting by enhancers. Here, we provide evidence that each core promoter element dynamically alters “enhancer responsiveness” by changing functional parameters of transcriptional bursting in developing *Drosophila* embryos. Quantitative live imaging analysis revealed that the timing and the continuity of burst induction are common regulatory steps on which core promoter elements impact. We also show that core promoters use the upstream TATA box as an amplifier of transcriptional bursting upon interaction with enhancers. Genome editing analysis of the pair-rule gene *fushi tarazu* further revealed that the endogenous TATA box and the downstream core promoter element are both essential for the correct expression and the function of key developmental genes in early embryos. We suggest that core promoter elements serve as a key regulatory module in converting enhancer activity into transcription dynamics during animal development.

## Introduction

Communication between enhancers and core promoters is critical for the temporal and spatial specificity of gene expression. Enhancers are distal regulatory elements that contain a cluster of binding sites for sequence-specific transcription factors and co-activators, which usually span several hundreds to thousands base pairs (bp) in length. Recent quantitative imaging studies reported that enhancers are responsible for driving transcriptional bursting from their target core promoters (e.g., Bartman et al. 2016; Fukaya et al. 2016; Larsson et al. 2019). However, little is known about the role of core promoters in the process. Core promoters are short segments of DNA, which typically range −40 to +40 relative to the +1 transcription start site (TSS). They serve as a docking site for general transcription factors and RNA polymerase II (Pol II) for the assembly of the pre-initiation complex. Importantly, core promoters alone are not sufficient to initiate productive transcription, but they require enhancer’s activation cues to assemble active transcription machineries. Thus, core promoters act as a “gateway to transcription”, converting enhancer activity into gene expression (reviewed in Vo Ngoc et al. 2019).

Core promoters consist of several sequence motifs located at fixed positions relative to the TSS. The TATA box (TATA) (Goldberg 1979) is typically located 25 to 30 bp upstream of the TSS and directly interacts with TATA-binding protein (TBP), a subunit of the transcription factor IID (TFIID) complex. Initiator (Inr), motif 10 element (MTE), and downstream core promoter element (DPE) are sequence motifs downstream of TATA (Smale and Baltimore 1989; Burke and Kadonaga 1996; Lim et al. 2004) and serve as an additional docking site for TFIID via direct interaction with TBP-associating factors (TAFs) (Burke and Kadonaga 1997; Chalkley and Verrijzer 1999; Theisen et al. 2010; Louder et al. 2016; Patel et al. 2018). In addition, core promoters often contain binding-sites for sequence-specific DNA binding proteins such as Zelda and GAGA factor (GAF) in *Drosophila* (Hendrix et al. 2008; Harrison et al. 2011; Chen et al. 2013). The precise composition of core promoter elements varies substantially among Pol II-transcribed genes (Ohler et al. 2002; FitzGerald et al. 2006; Chen et al. 2014), which is thought to play a key role in determining the level of gene expression by changing the responsiveness to enhancers (Ohtsuki et al. 1998; Butler and Kadonaga 2001; Zabidi et al. 2015; Arnold et al. 2017). However, it remains largely unclear how individual core promoter elements impact temporal dynamics of gene transcription in living multicellular organisms.

Importantly, quantitative measurement of enhancer responsiveness has been technically difficult with traditional bulk approaches because transcriptional output is controlled not only by core promoter sequences, but also by combinatory effects of surrounding regulatory landscapes such as enhancer strength and chromosome topology (reviewed in Furlong and Levine 2018). Moreover, there is no universal core promoter architecture, making it difficult to compare expression profiles of different genes. Here, we developed a live-imaging system that permits quantitative comparison of the roles of individual elements using a standardized, optimized synthetic core promoter placed under the control of a fixed enhancer. We have newly produced more than 15 *Drosophila* MS2 strains that contain a variety of core promoter modifications at different positions. Quantitative image analysis revealed that each core promoter elements dynamically alter the enhancer responsiveness by changing the functional parameters of transcriptional bursting in developing embryos. Our data suggest that the timing and the continuity of burst induction are common regulatory steps on which core promoter elements impact. We also show that core promoters use the upstream TATA to increase the amplitude of transcriptional bursting upon interaction with distal enhancers. Consistent with this, conversion of a natural TATA-less core promoter into a TATA-containing core promoter dramatically facilitated successive induction of strong bursts, supporting the idea that TATA serves as an “amplifier” of transcriptional bursting. In addition, promoter-proximal Zelda sites were found to exhibit a unique pioneering activity to facilitate burst induction at the beginning of new cell cycle, adding another layer of complexity in this process. To further dissect the function of core promoter elements in the context of endogenous genome, we combined genome editing and site-directed transgenesis. Using the pair-rule gene *fushi tarazu* (*ftz*), we show that both TATA and DPE mutations dramatically change transcription dynamics throughout the expression domains, resulting in disrupted spatial patterning of gene expression and misregulation of the downstream segment polarity gene *engrailed* (*en*). We suggest that core promoter elements serve as a key regulatory module in converting enhancer activity into transcription dynamics during animal development.

## Results

To quantitatively visualize how each core promoter element regulates enhancer responsiveness in living *Drosophila* embryos, we employed the MS2/MCP live-imaging method (Garcia et al. 2013; Lucas et al. 2013). We constructed a reporter system in which the *yellow* gene was placed under the control of the *Drosophila* synthetic core promoter (DSCP) (Pfeiffer et al. 2008), a modified *even-skipped* (*eve*) core promoter containing TATA, Inr, MTE, and DPE at optimal positions relative to the TSS (Figure 1A, Figure S1A). The use of a standardized core promoter architecture permits systematic comparison of individual elements in an unambiguous manner. A 24x MS2 RNA stem-loop sequence was engineered into the 5′ untranslated region (UTR) to enable visualization of nascent RNA production with a maternally provided MCP-GFP fusion protein. *DSCP-MS2-yellow* reporter gene was placed under the control of a full-length 1.5-kb *snail* (*sna*) shadow enhancer (Figure 1B), which drives expression in the ventral region of early embryos (Perry et al. 2010; Dunipace et al. 2011). In this synthetic locus, *sna* shadow enhancer is located ~6.5 kb away from the TSS, which is similar to the enhancer-promoter distance found at the endogenous *sna* locus (~7 kb). Transgenes were integrated into a specific genomic landing site via phiC31-mediated transgenesis (Groth et al. 2004; Venken et al. 2006).

**Figure 1.**
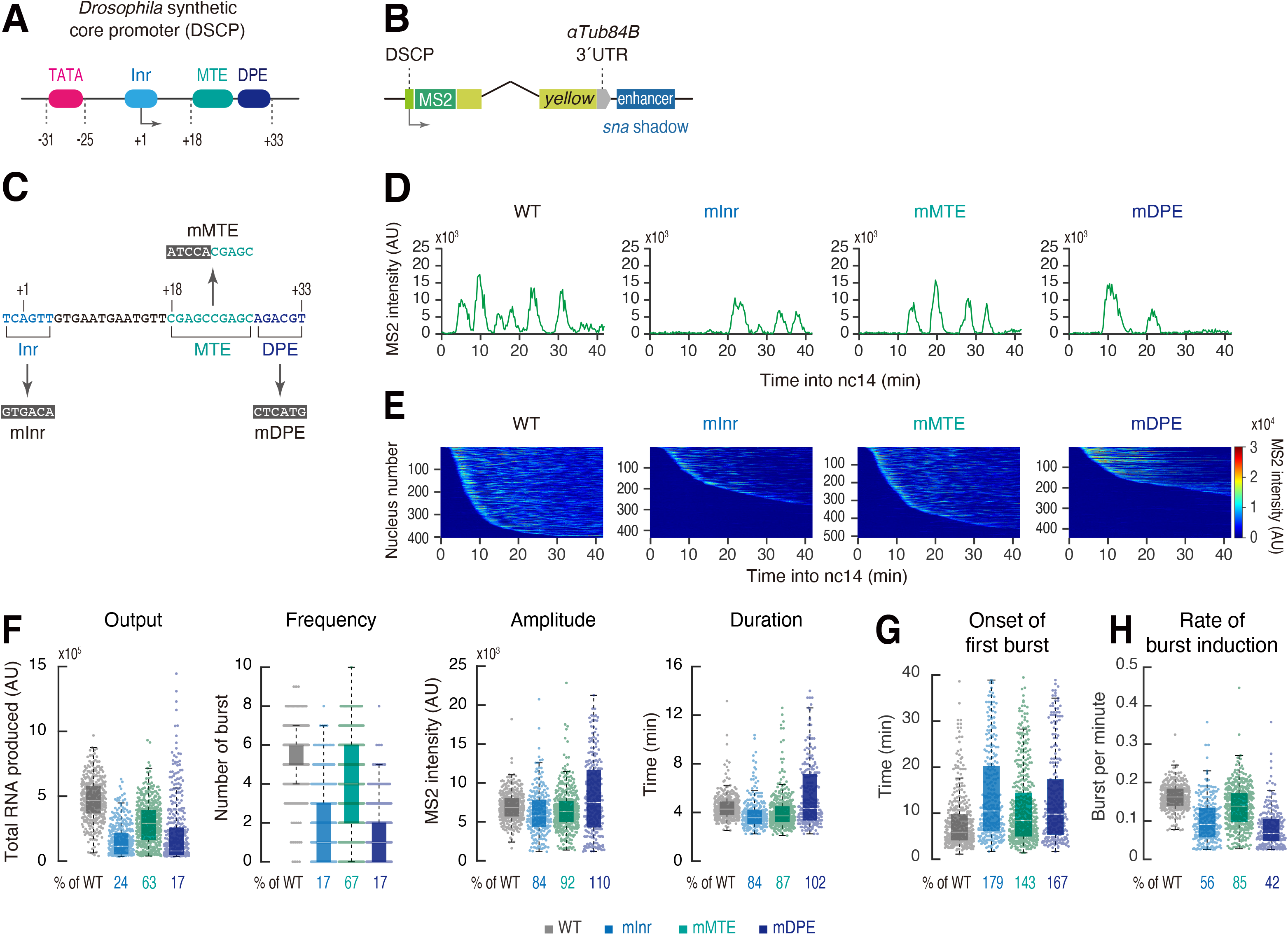
Downstream elements alter the frequency of transcriptional bursting. (A) Schematic representation of the *Drosophila* synthetic core promoter (DSCP). (B) Schematic representation of the *yellow* reporter gene containing the 155-bp DSCP, the 1.5-kb *sna* shadow enhancer, and 24x MS2 RNA stem loops (Bertrand et al. 1998) within the 5 ́ UTR. (C) Inr, MTE and DPE were mutated as indicated. (D) A representative trajectory of transcription activity of the MS2 reporter genes with WT (left), mInr (middle left), mMTE (middle right) and mDPE DSCP (right) in individual nuclei. AU; arbitrary unit. (E) MS2 trajectories for all analyzed nuclei. Each row represents the MS2 trajectory for a single nucleus. A total of 403, 435, 509 and 444 ventral-most nuclei, respectively, were analyzed from three independent embryos for the reporter genes with WT (left), mInr (middle left), mMTE (middle right) and mDPE DSCP (right). Nuclei were ordered by their onset of transcription in nc14. AU; arbitrary unit. (F) Boxplots showing the distribution of total output (left), burst frequency (middle left), burst amplitude (middle right) and burst duration (right). The box indicates the lower (25%) and upper (75%) quantile and the solid line indicates the median. Whiskers extend to the most extreme, non-outlier data points. A total of 403, 435, 509 and 444 ventral-most nuclei, respectively, were analyzed from three independent embryos for the reporter genes with WT, mInr, mMTE and mDPE DSCP. Median values relative to the WT reporter are shown at the bottom. AU; arbitrary unit. (G and H) Boxplots showing the distribution of the timing of first burst (G) and burst frequency normalized by the length of time after the first burst (H). The box indicates the lower (25%) and upper (75%) quantile and the solid line indicates the median. Whiskers extend to the most extreme, non-outlier data points. A total of 403, 435, 509 and 444 ventral-most nuclei, respectively, were analyzed from three independent embryos for the reporter genes with WT, mInr, mMTE and mDPE DSCP. Median values relative to the WT reporter are shown at the bottom.

First, we examined the downstream core promoter elements Inr, MTE and DPE by introducing mutations that compromise their direct interaction with TFIID (Figure 1C) (Burke and Kadonaga 1996; Burke and Kadonaga 1997; Kutach and Kadonaga 2000; Lim et al. 2004; Theisen et al. 2010). Notably, the mutation at the +18 to +22 positions eliminates MTE-dependent transcription without affecting DPE function (Lim et al. 2004; Theisen et al. 2010), allowing us to examine the individual contributions of MTE and DPE in the same core promoter context. Transcription activity was monitored from the entry into nuclear cycle 14 (nc14), when the major wave of zygotic genome activation occurs. To unambiguously compare the activities of different reporter genes at the same Dorsal-Ventral (DV) position, we focused on the ventral-most nuclei, as defined by the location of ventral furrow formation at the onset of gastrulation (Figure S1B). Quantitative image analysis revealed that the *sna* shadow enhancer induces fewer number of transcriptional bursts from the MTE mutant (mMTE) than the WT (Figure 1D, Movie S1). In comparison, even less frequent bursts were observed when the Inr mutant (mInr) and the DPE mutant (mDPE) were linked to the enhancer (Figure 1D, Movie S1). This trend was clearly seen when MS2 trajectories in all analyzed nuclei were visualized as a heat map (Figure 1E). These data suggest that downstream elements individually contribute to the induction of transcriptional bursting, with Inr/DPE playing a major role and MTE playing a more supplemental role. We then analyzed individual bursting events and quantified their functional parameters, including amplitude, duration, and frequency (Figure S1C). As seen in individual MS2 trajectories (Figure 1D), the frequency of transcriptional bursting and the total output were largely diminished upon mutation into downstream elements (Figure 1F, Figure S1D). In contrast, the amplitude and the duration remained less variable (Figure 1F). Importantly, all core promoter variants exhibited significant delay in the onset of the first round of transcription (Figure 1G), and substantial fraction of nuclei remained inactive during the analysis even though the *sna* shadow enhancer itself is in an active state as evidenced by the profile of the WT reporter (Figure S1E). We next determined rate of burst induction as a number of bursts divided by the time length after the onset of first burst for each active nucleus. There was an overall reduction in the efficiency of producing subsequent bursts after the first round of burst in all core promoter variants (Figure 1H). These data suggest that reduced bursting frequency is caused by overall reduction in the efficiency of initiating transcription in response to activation signals from distal enhancers. Essentially same results were observed when the *sna* shadow enhancer was replaced with another key developmental enhancer, *rhomboid* neuroectoderm element (*rho*NEE) (Ip et al. 1992) (Figure S2). We therefore suggest that the downstream elements support rapid and consecutive induction of transcriptional bursting, thereby facilitating production of a high level of nascent transcripts during early development.

We next examined the role of the upstream TATA, which we mutated according to previous characterization in order to abrogate its function (Figure 2A) (Butler and Kadonaga 2001; Lim et al. 2004). Live imaging analysis revealed that the TATA mutation causes more profound changes in overall transcription activity comparing to downstream modifications (Figure 2B-D, Figure S1F, Movie S2). Importantly, the *sna* shadow enhancer could only induce bursts with substantially lower amplitude from the TATA mutant (mTATA) (Figure 2D), suggesting that TATA mutation largely diminishes the number of Pol II released per burst. This result is consistent with recent single-cell RNA-seq studies that TATA affects the burst size in mammalian systems (Larsson et al. 2019; Ochiai et al. 2020). Burst duration was mostly unaffected by the loss of TATA (Figure 2D), suggesting that the size of each burst is mainly determined by the amplitude, but not the duration, of individual activation events. There also was a reduction in the frequency of transcriptional bursting when TATA was mutated (Figure 2D). As seen for downstream modifications (Figure 1G and H), reduced frequency appears to be attributed to delayed and inefficient burst induction (Figure 2E and F, Figure S1G). Strong TATA-dependency was also observed when the *rho*NEE was used for the analysis (Figure S2). We therefore suggest that core promoters use the upstream TATA to ensure rapid and consecutive induction of strong bursts upon interaction with enhancers.

**Figure 2.**
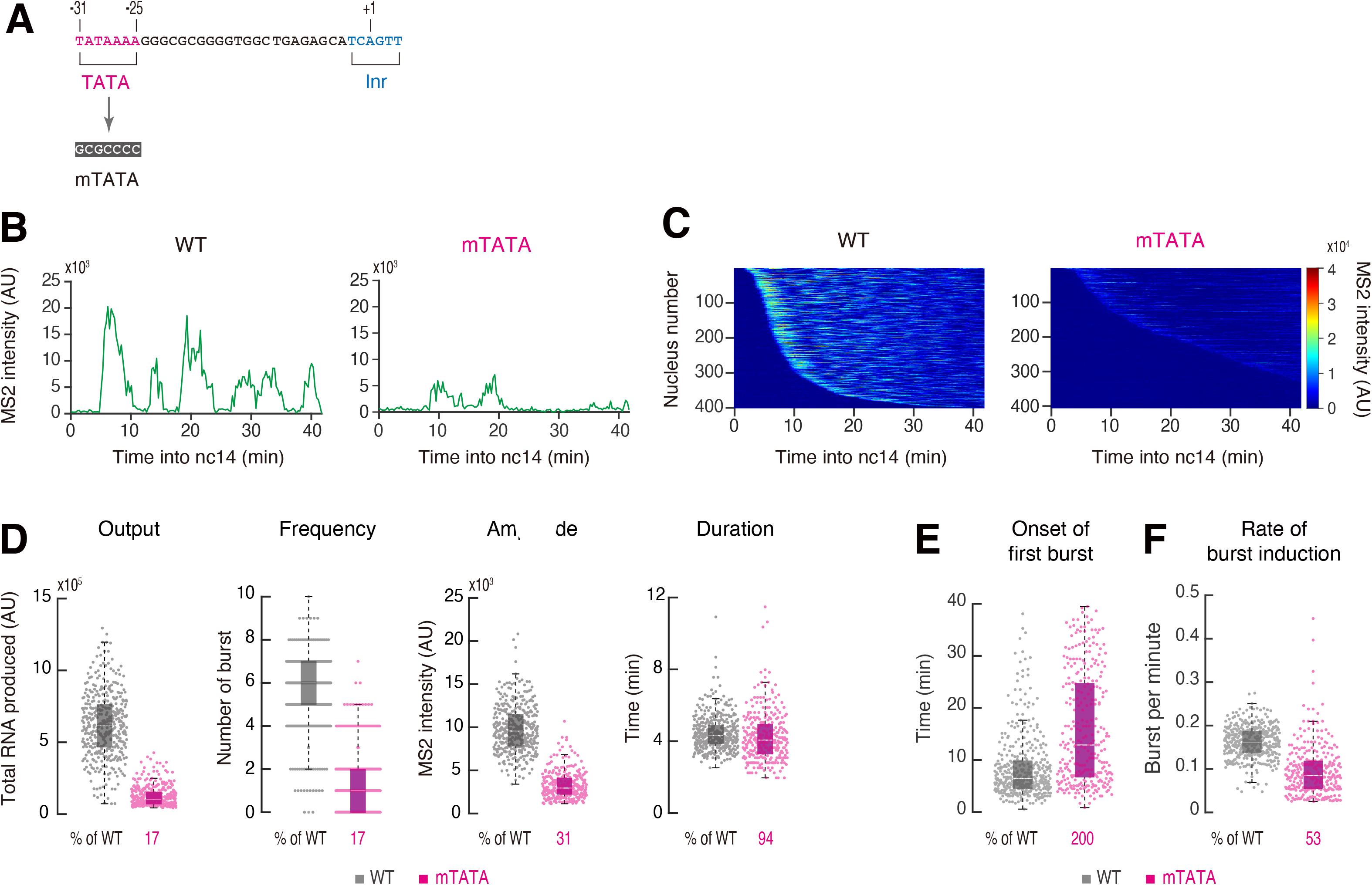
TATA acts as an amplifier of transcriptional bursting. (A) TATA was mutated as indicated. Modified core promoter was placed under the control of the *sna* shadow enhancer as illustrated in Figure 1B. (B) A representative trajectory of transcription activity of the MS2 reporter genes with WT (left) and mTATA DSCP (right) in individual nuclei. AU; arbitrary unit. (C) MS2 trajectories for all analyzed nuclei. Each row represents the MS2 trajectory for a single nucleus. A total of 401 and 406 ventral-most nuclei, respectively, were analyzed from three independent embryos for the reporter genes with WT (left) and mTATA DSCP (right). Nuclei were ordered by their onset of transcription in nc14. AU; arbitrary unit. (D) Boxplots showing the distribution of total output (left), burst frequency (middle left), burst amplitude (middle right) and burst duration (right). The box indicates the lower (25%) and upper (75%) quantile and the solid line indicates the median. Whiskers extend to the most extreme, non-outlier data points. A total of 401 and 406 ventral-most nuclei, respectively, were analyzed from three independent embryos for the reporter genes with WT and mTATA DSCP. Median values relative to the WT reporter are shown at the bottom. AU; arbitrary unit. (E and F) Boxplots showing the distribution of the timing of first burst (E) and burst frequency normalized by the length of time after the first burst (F). The box indicates the lower (25%) and upper (75%) quantile and the solid line indicates the median. Whiskers extend to the most extreme, non-outlier data points. A total of 401 and 406 ventral-most nuclei, respectively, were analyzed from three independent embryos for the reporter genes with WT and mTATA DSCP. Median values relative to the WT reporter are shown at the bottom.

We next tested if the conversion of a natural TATA-less core promoter into a TATA-containing core promoter has an opposite effect. It has been previously shown that core promoter regions of the *Drosophila* homeotic (Hox) genes are typically depleted of TATA (Juven-Gershon et al. 2008). Indeed, the core promoter of the Hox gene *labial* (*lab*) lacks an optimal TATA motif (Figure 3A and B). When the *lab* core promoter was placed under the control of the *sna* shadow enhancer, only weak and infrequent bursts were produced (Figure 3C and D). We then introduced two nucleotides substitution at the −28 and −30 positions to covert “TCTGAAA” to an optimal TATA motif, “TATAAAA” (Figure 3B). Intriguingly, this minimal modification dramatically increased overall transcription activities including the amplitude and the total output of transcriptional bursting (Figure 3C-E, Movie S3). In addition, the timing of first bursts (Figure 3F, Figure S3A) and the continuity of subsequent bursts (Figure 3G) were also augmented, resulting in a higher frequency of transcriptional bursting (Figure 3E). This result supports the idea that the upstream TATA serves an “amplifier” of transcriptional bursting. We suggest that endogenous TATA-less core promoters are naturally suboptimized to produce only an appropriate level of nascent transcripts upon interaction with enhancers by limiting the amplitude and the frequency of transcriptional bursting (see Discussion). Importantly, burst duration remained to be comparable even after this modification (Figure 3E), implicating that TATA does not affect the stability of active transcription machineries at the core promoter region.

**Figure 3.**
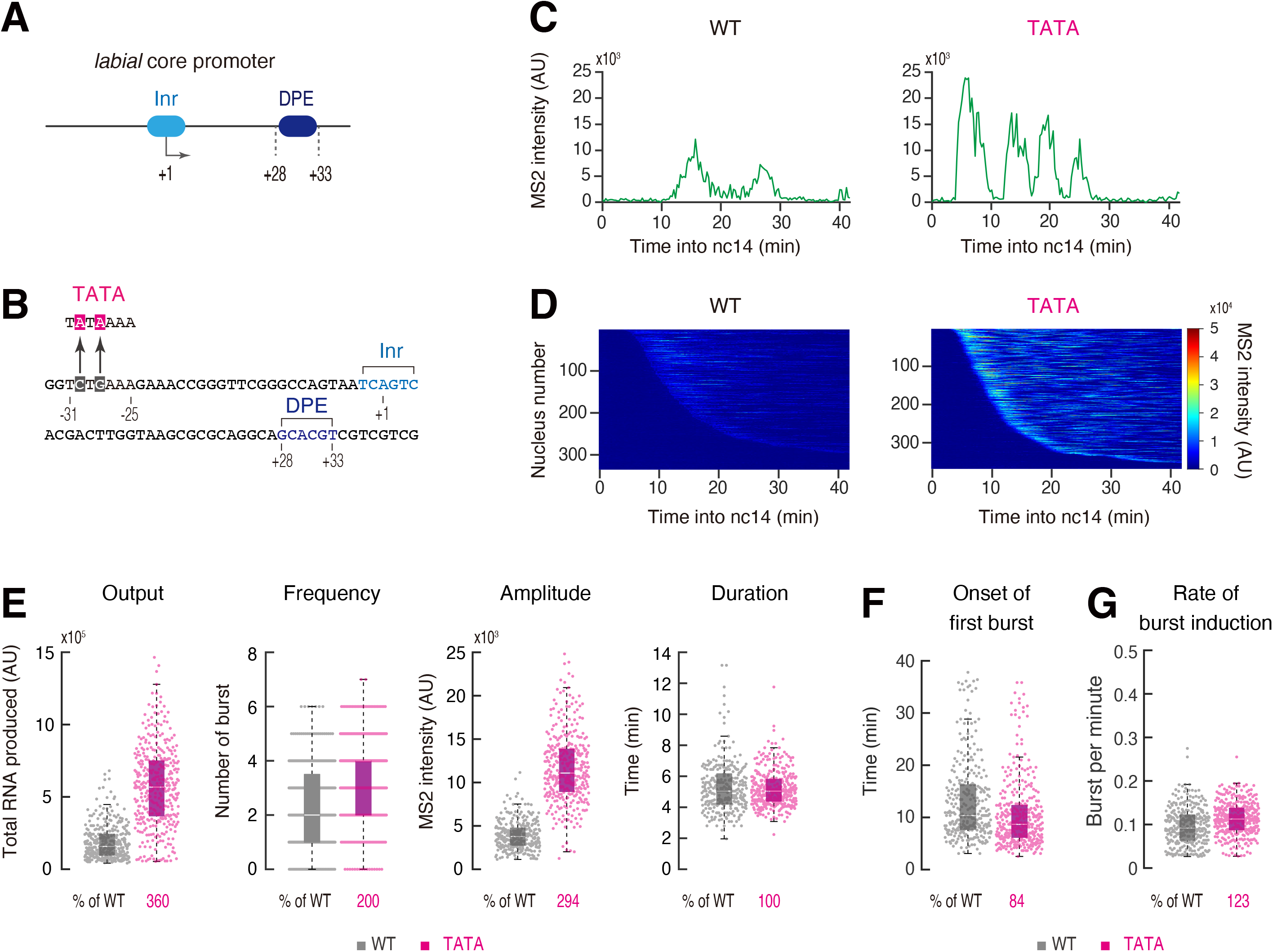
Endogenous TATA-less promoters are naturally suboptimized. (A) Endogenous *lab* core promoter contains Inr and DPE, but lacks TATA. The *sna* shadow enhancer was used for the analysis. (B) Two nucleotides substitution was introduced to contain an optimal TATA. (C) A representative trajectory of transcription activity of the MS2 reporter genes with unmodified (left) and modified *lab* core promoter (right) in individual nuclei. AU; arbitrary unit. (D) MS2 trajectories for all analyzed nuclei. Each row represents the MS2 trajectory for a single nucleus. A total of 336 and 371 ventral-most nuclei, respectively, were analyzed from three independent embryos for the reporter genes with unmodified (left) and modified *lab* core promoter (right). Nuclei were ordered by their onset of transcription in nc14. AU; arbitrary unit. (E) Boxplots showing the distribution of total output (left), burst frequency (middle left), burst amplitude (middle right) and burst duration (right). The box indicates the lower (25%) and upper (75%) quantile and the solid line indicates the median. Whiskers extend to the most extreme, non-outlier data points. A total of 336 and 371 ventral-most nuclei, respectively, were analyzed from three independent embryos for the reporter genes with unmodified and modified *lab* core promoter. Median values relative to the unmodified *lab* core promoter are shown at the bottom. AU; arbitrary unit. (F and G) Boxplots showing the distribution of the timing of first burst (F) and burst frequency normalized by the length of time after the first burst (G). The box indicates the lower (25%) and upper (75%) quantile and the solid line indicates the median. Whiskers extend to the most extreme, non-outlier data points. A total of 336 and 371 ventral-most nuclei, respectively, were analyzed from three independent embryos for the reporter genes with unmodified and modified *lab* core promoter. Median values relative to the unmodified *lab* core promoter are shown at the bottom.

In *Drosophila*, core promoters often contain binding sites for sequence-specific DNA binding proteins, Zelda and GAF (Lee et al. 2008; Harrison et al. 2011), that are thought to recruit chromatin remodeling complexes such as NURF to increase chromatin accessibility of regulatory regions (Tsukiyama and Wu 1995; Fuda et al. 2015; Sun et al. 2015). However, it remains unclear how they modulate core promoter functions because previous studies have mainly focused on their roles at enhancer regions (e.g., Foo et al. 2014; Sun et al. 2015; Dufourt et al. 2018; Yamada et al. 2019). DSCP contains a GAF-binding site (GAGA) from the *eve* core promoter (Figure S1A) (Ohtsuki and Levine 1998), but lacks any known Zelda binding motifs. To examine how promoter-proximal GAGA and Zelda sites influence transcription dynamics, we either mutagenized GAGA site or added Zelda sites (Figure 4A). Bursting activities were moderately diminished upon GAGA mutation (Figure 4B-D). On the other hand, addition of Zelda sites led to rapid induction of strong bursts (Figure 4B and C, Figure S3B) and an overall increase in transcription activities (Figure 4D, Movie S4). Intriguingly, while addition of Zelda sites slightly increased the continuity of burst induction after the first round of transcription (Figure 4F), the timing of initial burst was found to be dramatically accelerated (Figure 4E, Figure S3B). Consistent with this, the level of RNA production during the first 20 min of the analysis was more strongly increased by the addition of Zelda sites than the rest of 20 min (Figure S3C). Such a time-dependent activity was not clearly seen for other core promoter elements (Figure S3D and E), suggesting that promoter-proximal Zelda sites have a unique pioneering activity that increases the enhancer responsiveness especially at the onset of nc14, when the major wave of zygotic genome activation starts to take place.

**Figure 4.**
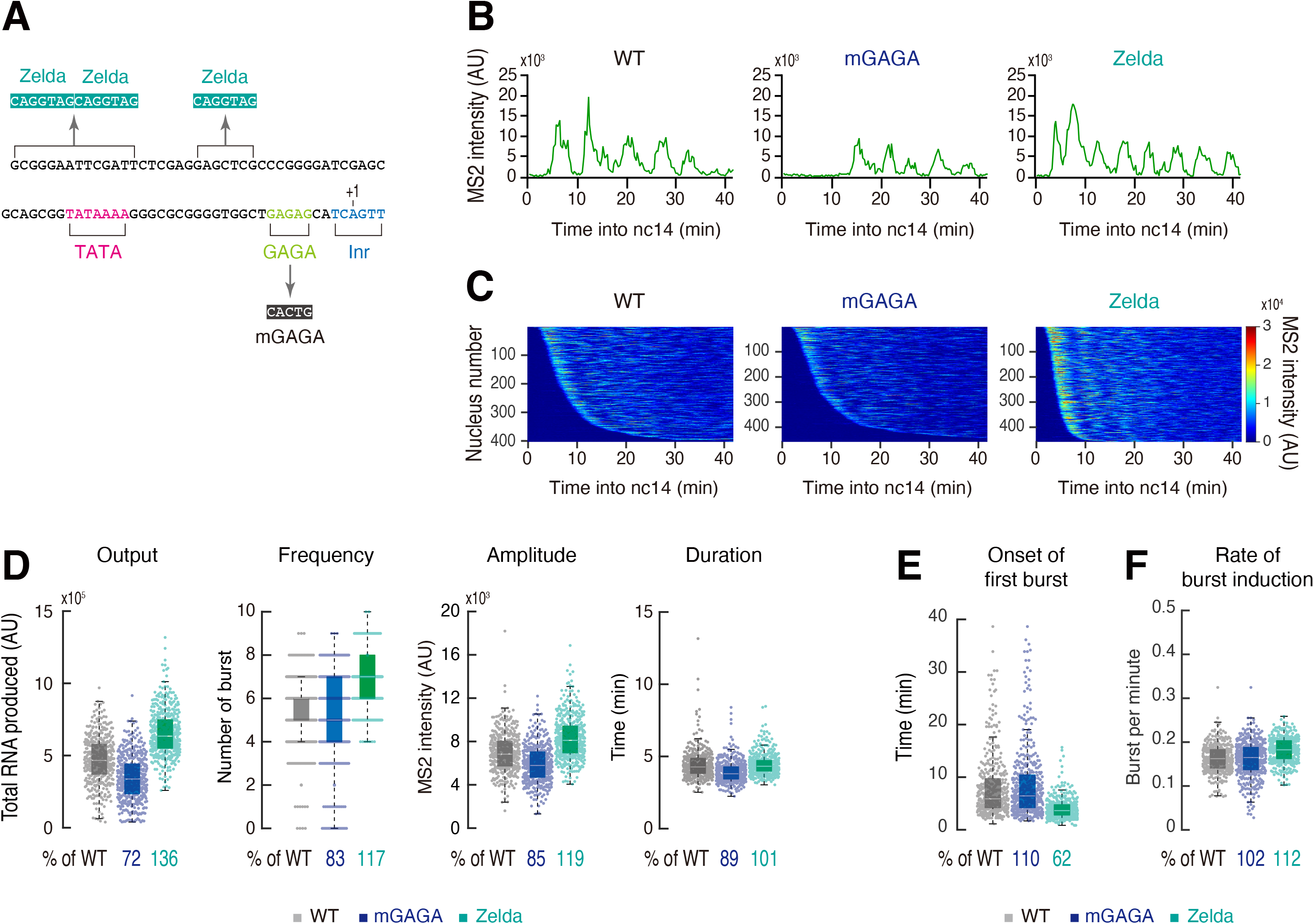
Promoter-proximal Zelda exhibits a unique pioneering activity. (A) DSCP was modified to mutate GAGA site or add three optimal Zelda binding sites. The *sna* shadow enhancer was used for the analysis. (B) A representative trajectory of transcription activity of the MS2 reporter genes with WT (left), mGAGA (middle) and Zelda DSCP (right) in individual nuclei. AU; arbitrary unit. (C) MS2 trajectories for all analyzed nuclei. Each row represents the MS2 trajectory for a single nucleus. A total of 403, 458 and 458 most ventral-nuclei, respectively, were analyzed from three individual embryos for the reporter genes with WT (left), mGAGA (middle) and Zelda DSCP (right). Nuclei were ordered by their onset of transcription in nc14. Panel of WT is the same as the panel in Figure 1E. AU; arbitrary unit. (D) Boxplots showing the distribution of total output (left), burst frequency (middle left), burst amplitude (middle right) and burst duration (right). The box indicates the lower (25%) and upper (75%) quantile and the solid line indicates the median. Whiskers extend to the most extreme, non-outlier data points. A total of 403, 458 and 458 most ventral-nuclei, respectively, were analyzed from three individual embryos for the reporter genes with WT, mGAGA and Zelda DSCP. Median values relative to the WT DSCP reporter are shown at the bottom. Plot of WT is the same as the plot in Figure 1F. AU; arbitrary unit. (E and F) Boxplots showing the distribution of the timing of first burst (E) and burst frequency normalized by the length of time after the first burst (F). The box indicates the lower (25%) and upper (75%) quantile and the solid line indicates the median. Whiskers extend to the most extreme, non-outlier data points. A total of 403, 458 and 458 ventral-most nuclei, respectively, were analyzed from three individual embryos for the reporter genes with WT, mGAGA and Zelda DSCP. Median values relative to the WT reporter are shown at the bottom. Plot of WT is the same as the plot in Figure 1G and H.

To explore how core promoter elements impact enhancer-promoter interactions at the endogenous locus, we next focused on one of the best studied developmental patterning genes, *fushi tarazu* (*ftz*) (Wakimoto and Kaufman 1981; Nüsslein-Volhard 1982; Hafen et al. 1984; Kuroiwa et al. 1984; Wakimoto et al. 1984). *ftz* is expressed in seven transverse stripes spanning across the anterior-posterior (AP) axis and is regulated by multiple enhancers located 5′ and 3′ of the transcription unit (Hiromi et al. 1985; Hiromi and Gehring 1987; Pick et al. 1990; Calhoun and Levine 2003; Schroeder et al. 2011). The *ftz* core promoter contains TATA, Inr and DPE elements (Figure 5A) (Juven-Gershon et al. 2008). Reporter assays in *Drosophila* S2 cultured cells have suggested that DPE, but not TATA, is specifically required for the activation of the *ftz* core promoter by the homeodomain-containing transcription factor Caudal (Juven-Gershon et al. 2008). In this model, mutation of DPE is expected to most severely affect *ftz* expression in stripe 5 and 6 because they are regulated by Caudal-dependent enhancers (Macdonald and Struhl 1986). To test this idea, we developed a genome engineering approach for visualizing impacts of core promoter modification at the endogenous locus. First, the entire *ftz* transcription unit was replaced with attP site via CRISPR/Cas9-mediated genome editing. Subsequently, a full-length *ftz* transcription unit containing the modified core promoter and 24x MS2 sequence was integrated into the attP site via phiC31-mediated transgenesis (Figure 5B, Figure S4A). Fluorescent *in situ* hybridization assay revealed that *ftz* expression is diminished equally across all seven stripes upon DPE mutation (Figure 5C), indicating that the *ftz* core promoter uses DPE for responding not only to Caudal-dependent stripe 5 and 6 enhancers but also to all the other stripe enhancers. Moreover, *ftz* expression was almost completely lost upon TATA mutation (Figure 5C). As a consequence of irregular *ftz* patterning, expression of the downstream segment polarity gene *engrailed* (*en*) was lost from even-numbered stripes in mTATA and mDPE embryos (Figure 5D). These results suggest that both TATA and DPE are required for the correct expression and the function of *ftz* during embryogenesis.

**Figure 5.**
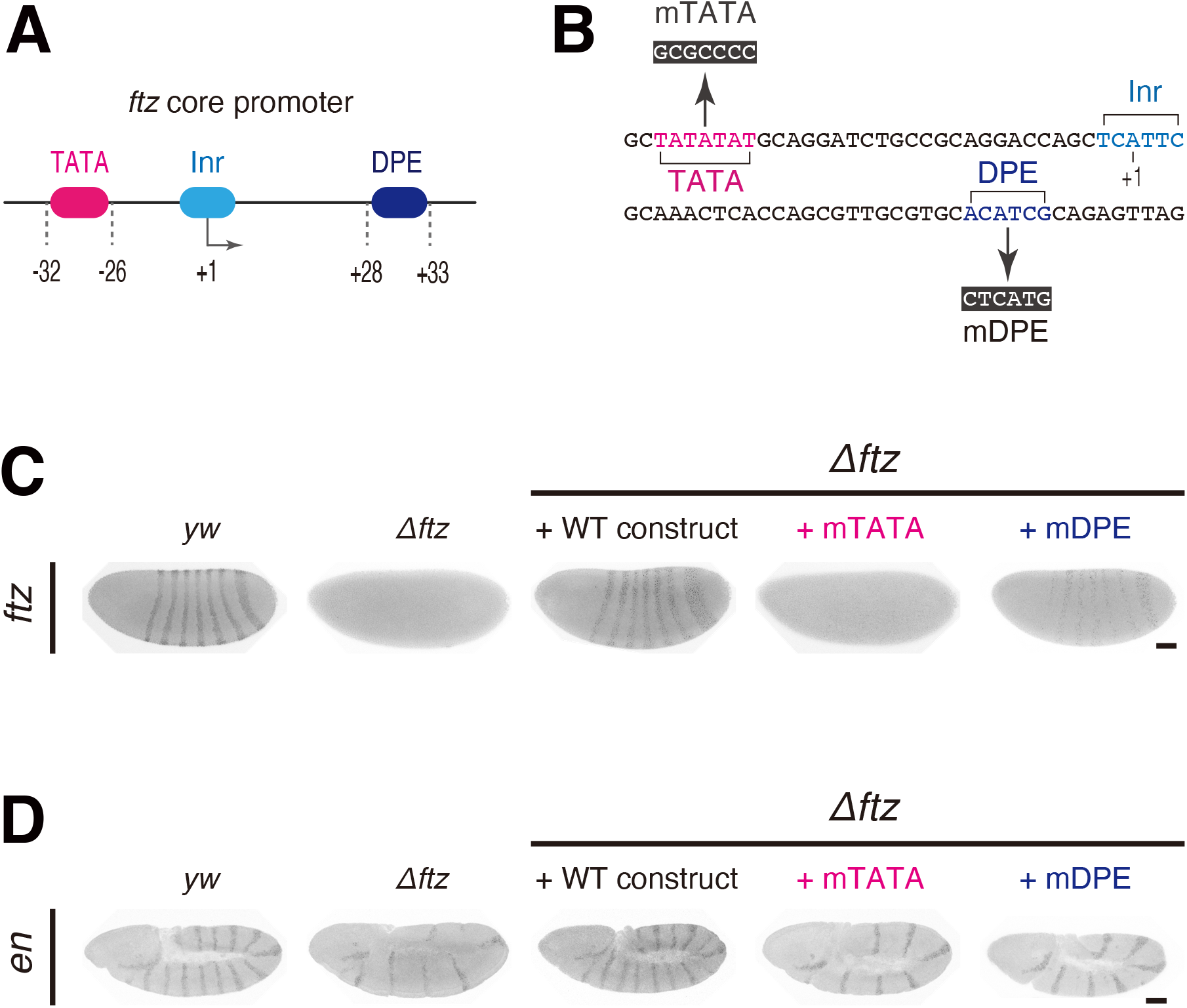
TATA and DPE are both required for proper *ftz* expression. (A) *ftz* core promoter contains TATA, Inr and DPE. (B) TATA and DPE were mutated as indicated. (C) Fluorescent *in situ* hybridization of *ftz*. Embryos at late nc14 are shown. *yw* embryo is shown as a control. *ftz-MS2* constructs were integrated into the attP site at the *Δftz* allele. Images are cropped and rotated to align embryos (anterior to the left and posterior to the right). Scale bar indicates 50 μm. (D) Fluorescent *in situ* hybridization of *en*. Embryos after germband extension are shown. *yw* embryo is shown as a control. Images are cropped and rotated to align embryos (anterior to the left and posterior to the right). Brightness of each embryo was differentially adjusted for visualization of *en* expression pattern. Scale bar indicates 50 μm.

To elucidate the molecular basis underlying these phenotypes, we carried out live-imaging analysis of individual *ftz-MS2* alleles. We focused on anterior stripe 1/2 and posterior stripe 5/6 (Figure 6A, Movie S5) because they are known to be regulated by different enhancers located 5′ and 3′ of the gene (Figure S4A) (Calhoun and Levine 2003; Schroeder et al. 2011). While both the mTATA and mDPE *ftz* alleles failed to restore *en* expression (Figure 5D), their MS2 profiles differ dramatically. The mDPE allele produced delayed discontinuous but strong bursting activity (Figure 6B and C), whereas the mTATA allele further led to the overall diminishment of MS2 intensity in all analyzed stripes (Figure 6B and C). As a consequence, the mTATA allele exhibited a ~82-89% reduction in the total RNA production, while the mDPE allele showed a more modest reduction (Figure 6D). We then characterized the functional parameters of MS2 trajectories. As we previously reported (Yokoshi et al. 2020), individual bursting events were hard to be discerned due to the continuity of bursting activities (Figure 6B), suggesting that the endogenous genomic configurations are highly optimized for efficient production of transcriptional bursting within a short period of time. To minimize ambiguity in defining individual bursting events, we quantified the instantaneous fraction of active nuclei (Figure 6E) and the mean MS2 intensity in actively transcribing nuclei (Figure 6F). We found that mutation of TATA and DPE both diminish the fraction of active nuclei (Figure 6E) and also delay the onset of transcription in all stripe regions (Figure S4B and C). Importantly, the level of MS2 intensity was more severely reduced in the mTATA allele (Figure 6F), suggesting that the TATA mutation reduces the number of Pol II entering into productive elongation even when core promoter in an active state. These profiles were similar to those of mDPE and mTATA DSCP linked to the *sna* shadow enhancer at the synthetic locus (Figure S4D and E). We therefore suggest that DPE and TATA differentially regulate the enhancer responsiveness by changing the efficiency and the strength of burst induction. Lastly, to determine how misregulation of bursting activities affects the stripe formation, we computationally reconstituted the spatial patterning of *ftz* by calculating the total RNA production at each nucleus and the *ftz* mRNA turnover rate in early embryos (half-life: 7 min) (Edgar et al. 1986). Although the mDPE allele exhibited somewhat stronger MS2 activities than the mTATA (Figure 6B-F), both resulted in highly sporadic stripe patterns (Figure S5). Thus, we concluded that the endogenous configuration of *ftz* core promoter is essential for the uniform expression at the stripe regions to ensure proper control of downstream genes such as *en* during embryo segmentation.

**Figure 6.**
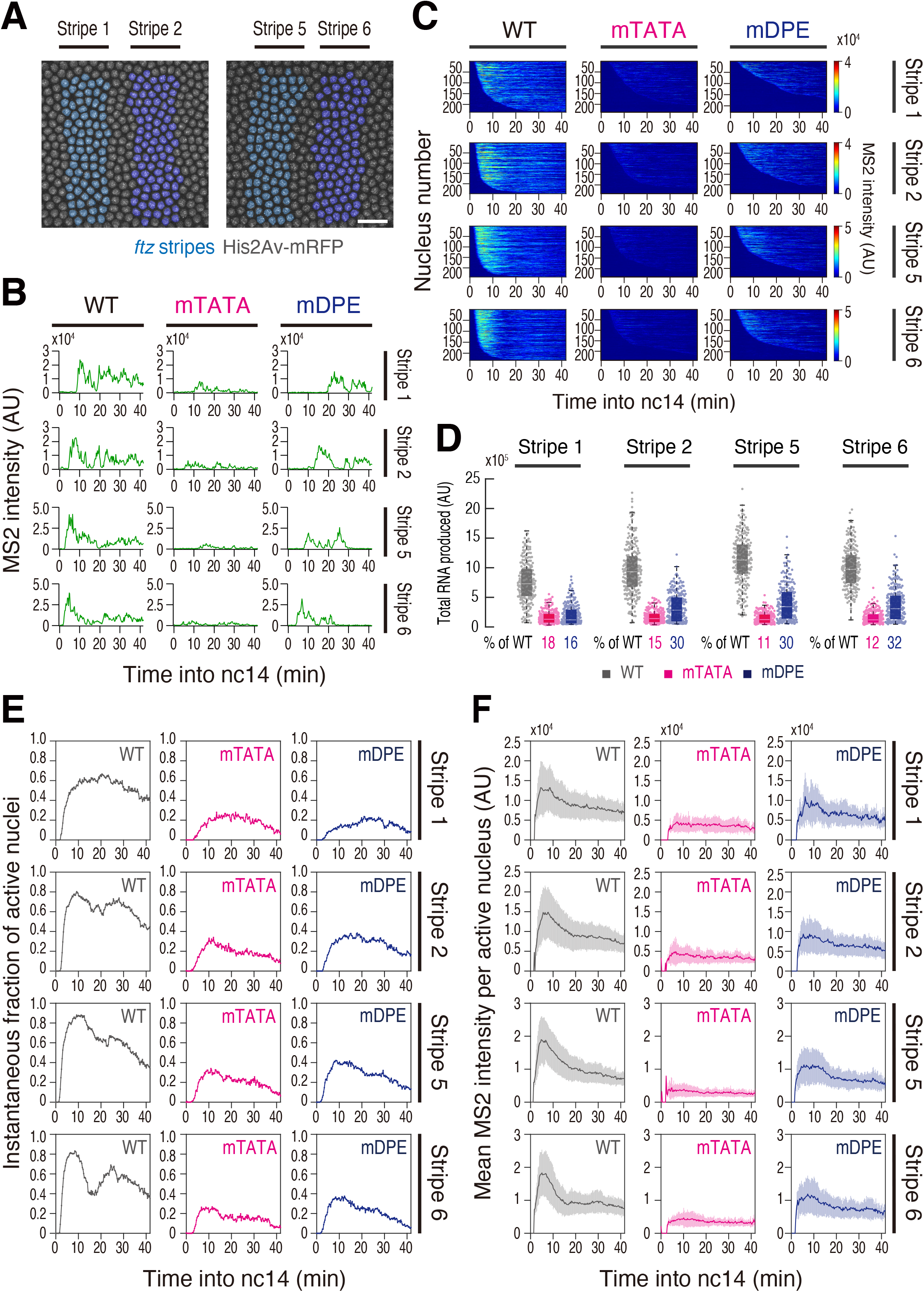
TATA and DPE differentially regulate *ftz* transcription. (A, left) False-coloring of nuclei at stripe 1 (cyan) and stripe 2 (blue). (A, right) False-coloring of nuclei at stripe 5 (cyan) and stripe 6 (blue). The maximum projected image of a histone marker (His2Av-mRFP) is shown in gray. Images are oriented with anterior to the left. Scale bar indicates 20 μm. (B) A representative trajectory of transcription activity of *ftz-MS2* in individual nuclei. AU; arbitrary unit. (C) MS2 trajectories for all analyzed nuclei. Each row represents the MS2 trajectory for a single nucleus. A total of 234, 236 and 228 nuclei at stripe 1, 248, 241 and 242 nuclei at stripe 2, 243,238 and 240 nuclei at stripe 5, and 232, 243 and 240 nuclei at stripe 6 were analyzed from three individual embryos for the *ftz-MS2* with WT, mTATA and mDPE core promoter, respectively. Nuclei were ordered by their onset of transcription in nc14. AU; arbitrary unit. (D) Boxplot showing the distribution of total output. The box indicates the lower (25%) and upper (75%) quantile and the solid line indicates the median. Whiskers extend to the most extreme, non-outlier data points. A total of 234, 236 and 228 nuclei at stripe 1, 248, 241 and 242 nuclei at stripe 2, 243,238 and 240 nuclei at stripe 5, and 232, 243 and 240 nuclei at stripe 6 were analyzed from three individual embryos for the *ftz-MS2* with WT, mTATA and mDPE core promoter, respectively. Median values relative to the *ftz-MS2* with WT core promoter are shown at the bottom. AU; arbitrary unit. (E) Instantaneous fraction of actively transcribing nuclei at each expression domain. A total of 234, 236 and 228 nuclei at stripe 1, 248, 241 and 242 nuclei at stripe 2, 243,238 and 240 nuclei at stripe 5, and 232, 243 and 240 nuclei at stripe 6 were analyzed from three individual embryos for the *ftz-MS2* with WT, mTATA and mDPE core promoter, respectively. (F) Mean MS2 intensity per actively transcribing nucleus at each expression domain. A total of 234, 236 and 228 nuclei at stripe 1, 248, 241 and 242 nuclei at stripe 2, 243,238 and 240 nuclei at stripe 5, and 232, 243 and 240 nuclei at stripe 6 were analyzed from three individual embryos for the *ftz-MS2* with WT, mTATA and mDPE core promoter, respectively. Error bars represent the standard deviation of the mean across active nuclei at given time.

## Discussion

In this study, we provided evidence that each core promoter element differentially modulates the responsiveness to enhancers in early *Drosophila* embryos. Our data suggest that the downstream elements, Inr, MTE and DPE mainly affect the frequency of transcriptional bursting by changing the timing of first burst and the continuity of subsequent bursts (Figure 1). On the other hand, TATA mutation dramatically diminished overall transcription activity including the bursting amplitude (Figure 2 and 3). Our genome engineering approach further revealed that the endogenous *ftz* core promoter requires both TATA and DPE to initiate rapid and productive transcription upon interaction with stripe enhancers (Figure 5 and 6). Importantly, recent cryo-EM studies have revealed that TFIID recognizes the core promoter in a stepwise fashion (Cianfrocco et al. 2013; Louder et al. 2016; Patel et al. 2018), with initial contact with Inr, MTE and DPE through TAF1/2 subunits, followed by dynamic structural rearrangement that facilitates recognition of the upstream TATA by TBP. We speculate that the rate of initial TFIID loading onto the downstream core promoter region mainly affects the timing and the frequency, and the subsequent TBP loading onto the upstream TATA helps to further increase the bursting amplitude. Supporting this view, a recent biochemical study suggested that downstream core promoter interactions of TFIID increase the efficiency of transcription reinitiation in yeast (Joo et al. 2017). Addition of Zelda sites was found to augment both of these parameters (Figure 4), suggesting that the promoter-opening by Zelda facilitates the initial TFIID loading and the subsequent TBP loading during the onset of nc14. It has been previously shown that Zelda is pre-loaded onto a thousand of promoter regions prior to zygotic genome activation in early *Drosophila* embryos (Harrison et al. 2011). Thus, it appears that promoter-proximal Zelda sites increase the responsiveness to enhancers by making core promoters poised for activation even before distal enhancers start to drive transcription. We suggest that these multi-layered mechanisms help to diversify the enhancer responsiveness of core promoters across the genome because there are substantial variations in the composition of core promoter elements among Pol II-transcribed genes. For example, it is estimated that only a small fraction of protein coding genes contain TATA, both in human and *Drosophila* (Ohler et al. 2002; FitzGerald et al. 2006; Chen et al. 2014). Importantly, our data showed that the TATA-less core promoter of the Hox gene *lab* is naturally suboptimized to limit the level of total RNA production by reducing the amplitude of transcriptional bursting (Figure 3). As suboptimal transcription factor binding sites are important for tissue-specific gene activation by enhancers (Crocker et al. 2015; Farley et al. 2015), suboptimal core promoter architectures might play a key role for ensuring the spatial and temporal specificity of gene expression during animal development. Alternatively, it can be possible that a subset of enhancers are capable of driving strong bursts even when TATA or DPE is not present as suggested by previous enhancer trapping assay in *Drosophila* (Butler and Kadonaga 2001). Our study will serve as a critical starting point toward understanding of how temporal dynamics of gene expression is encoded in the eukaryotic genome.

## Acknowledgment

We thank Bomyi Lim and Tyler Heist for sharing nuclei segmentation and tracking code, Hitomi Takishita and Misako Sato for fly husbandry and the Bloomington *Drosophila* Stock Center for fly strains. We are also grateful to Michael Levine, Tyler Heist, Yuko Hasegawa, Life Science Editors and members of the Fukaya laboratory for critical comments on the manuscript. M.Y is supported by the Grant-in-Aid for Early-Career Scientists (20K15710) from the Japan Society for the Promotion of Science, and the Employment Stability Support for Young Researchers from the University of Tokyo. M.C is supported by the FPI research grant (FPI2015/074837) from the MINECO-Feder, and the FisyMat PhD Student Research program from University of Granada. This study was funded by the Grant-in-Aid for Scientific Research on Innovative Areas (Research in a Proposed Research Area) (20H05357), the Grants-in-Aid for Scientific Research (B) (19H03154), the Grants-in-Aid for Challenging Research (Exploratory) (19K22378), the Grants-in-Aid for Research Activity Start-up (18H06040) from the Japan Society for the Promotion of Science, the Grants-in-Aid for Leading Initiative for Excellent Young Researchers from the Ministry of Education, Culture, Sports, Science and Technology in Japan, Tomizawa Jun-ichi & Keiko Fund of Molecular Biology Society of Japan for Young Scientist, research grants from the Mochida Memorial Foundation for Medical and Pharmaceutical Research, the Nakajima Foundation, the Inamori Foundation, the Takeda Science Foundation, the Sumitomo Foundation and the Senri Life Science Foundation.

## Author Contributions

M.Y performed the experiments. M.Y and T.F analyzed the data. M.C developed image analysis code. T.F wrote the manuscript. All the authors discussed the results and approved the manuscript.

## Materials and Methods

### Experimental model

In all live-imaging experiments, we studied *Drosophila melanogaster* embryos at nuclear cycle 14. The following fly lines were used in this study: *nanos>MCP-GFP, His2Av-mRFP/CyO* (Yokoshi et al. 2020), *DSCP_WT_-MS2-yellow-sna shadow enhancer* (Yokoshi et al. 2020), *DSCP_mTATA_-MS2-yellow-sna shadow enhancer* (this study), *DSCP_mInr_-MS2-yellow-sna shadow enhancer* (this study), *DSCP_mMTE_-MS2-yellow-sna shadow enhancer* (this study), *DSCP_mDPE_-MS2-yellow-sna shadow enhancer* (this study), *DSCP_mGAGA_-MS2-yellow-sna shadow enhancer* (this study), *DSCP_3xZelda_-MS2-yellow-sna shadow enhancer* (this study), *DSCP_WT_-MS2-yellow-rhoNEE* (this study), *DSCP_mTATA_-MS2-yellow-rhoNEE* (this study), *DSCP_mInr_-MS2-yellow-rhoNEE* (this study), *DSCP_mMTE_-MS2-yellow-rhoNEE* (this study), *DSCP_mDPE_-MS2-yellow-rhoNEE* (this study), *lab_WT_-MS2-yellow-sna shadow enhancer* (this study), *lab_TATA_-MS2-yellow-sna shadow enhancer* (this study), *Δftz-attP/TM6* (this study), *ftz WT core promoter-ftz-HA-24xMS2* (this study), *ftz mTATA core promoter-ftz-HA-24xMS2/TM6* (this study), and *ftz mDPE core promoter-ftz-HA-24xMS2/TM6* (this study).

### Site specific transgenesis by phiC31 system

All reporter plasmids were integrated into a unique landing site on the third chromosome using VK00033 strain (Venken et al. 2006). PhiC31 was maternally provided using vas-phiC31 strain (Bischof et al. 2007). Microinjection was performed as previously described (Ringrose 2009). In brief, 0-1 h embryos were collected and dechorionated with bleach. Aligned embryos were dried with silica gel for ~7 min and covered with FL-100-1000CS silicone oil (Shin-Etsu Silicone). Subsequently, microinjection was performed using FemtoJet (Eppendorf) and DM IL LED inverted microscope (Leica) equipped with M-152 Micromanipulator (Narishige). Injection mixture typically contains ~500 ng/μl plasmid DNA, 5 mM KCl, 0.1 mM phosphate buffer, pH 6.8. mini-white marker was used for screening.

### Core promoter modification at endogenous locus

First, endogenous *ftz* transcription unit was removed and replaced with attP site by CRISPR/Cas9-mediated genome editing. Two pCFD3 gRNA expression plasmids and pBS-attP-dsRed donor plasmid were co-injected using *nanos-Cas9/CyO* strain (Ren et al. 2013). Injection mixture contains 500 ng/μl pCFD3 gRNA expression plasmids, 500 ng/μl pBS-attP-dsRed donor plasmid, 5 mM KCl, 0.1 mM phosphate buffer, pH 6.8. Resulting *ftz* allele were balanced over TM6. Subsequently, *ftz-MS2* plasmid was integrated into the attP site using *Δftz-attP/TM6* strain. Corresponding MS2 plasmid and p3×3-EGFP.vas-int.NLS plasmid (addgene #60948) were co-injected. Injection mixture contains 500 ng/μl MS2 plasmid, 500 ng/μl phiC31 expression plasmid, 5 mM KCl, 0.1 mM phosphate buffer, pH 6.8. mini-white marker was used for screening. Resulting *ftz* complementation alleles were balanced over TM6. After phiC31-mediated integration, extra sequences derived from plasmid backbone, ampicillin resistant gene and mini-white marker gene were also incorporated into the *ftz* locus.

### cDNA synthesis

Total RNA was extracted from 40 adults of Oregon-R using TRIzol reagent (Thermo Fisher) followed by chloroform purification and isopropanol precipitation. Three μg of total RNA was subjected to reverse transcription using PrimeScript 1st strand cDNA Synthesis Kit (Takara).

### Preparation of probes for *in situ* hybridization

Antisense RNA probes labeled with digoxigenin (DIG RNA Labeling Mix 10 × conc, Roche) and biotin (Biotin RNA Labeling Mix 10 × conc, Roche) were *in vitro* transcribed using *in vitro* Transcription T7 Kit (Takara). Template DNA for *ftz* probe was PCR amplified from genomic DNA using primers (5 ́-CGT AAT ACG ACT CAC TAT AGG GTG GGG AAG AGA GTA ACT GAG CAT CGC-3 ́) and (5 ́-ATT CGC AAA CTC ACC AGC GT-3 ́). Template DNA for *en* probe was PCR amplified from cDNA using primers (5 ́-CGT AAT ACG ACT CAC TAT AGG GCA TGA ACT TGC TTT AGC ACA AAC ATT TCG-3 ́) and (5 ́-CAA CTA ATT CAG TCG TTG CGC TCG-3 ́).

### Fluorescence *in situ* hybridization

Embryos were dechorionated and fixed in fixation buffer (1 ml of 5x PBS, 4 ml of 37% formaldehyde and 5 ml of Heptane) for ~25 min at room temperature. Vitelline membrane was then removed by shaking embryos in a biphasic mixture of heptane and methanol for ~1 min. Antisense RNA probes labeled with digoxigenin and biotin were used. Hybridization was performed at 55 ̊C overnight in hybridization buffer (50% formamide, 5x SSC, 50 μg/ml Heparin, 100 μg/ml salmon sperm DNA, 0.1% Tween-20). Subsequently, embryos were washed with hybridization buffer at 55 ̊C and incubated with Western Blocking Reagent (Roche) at room temperature for one hour. Then, embryos were incubated with sheep anti-digoxigenin (Roche) and mouse anti-biotin primary antibodies (Invitrogen) at 4 ̊C for overnight, followed by incubation with Alexa Fluor 555 donkey anti-sheep (Invitrogen) and Alexa Flour 488 goat anti-mouse (Invitrogen) fluorescent secondary antibodies at room temperature for one hour. DNA was stained with DAPI, and embryos were mounted in ProLong Gold Antifade Mountant (Thermo Fisher Scientific). Imaging was performed on a Zeiss LSM 900 confocal microscope. Plan-Apochromat 20x / 0.8 N.A. objective was used. Images were captured in 16-bit. Maximum projections were obtained for all z-sections, and resulting images were shown. Brightness of images was linearly adjusted using Fiji (https://fiji.sc).

### MS2 live imaging

Virgin females of *nanos>MCP-GFP, His2Av-mRFP/CyO* (Yokoshi et al. 2020) were mated with males carrying the MS2 allele. The resulting embryos were dechorionated and mounted between a polyethylene membrane (Ube Film) and a coverslip (18 mm × 18 mm), and embedded in FL-100-450CS (Shin-Etsu Silicone). Embryos were imaged using a Zeiss LSM 800 (Figure 1 and 4) or LSM900 (Figure 2, 3, 6 and S2). Temperature was kept in between 23.5 to 25.0 ̊C during imaging. Plan-Apochromat 40x / 1.4 N.A. oil immersion objective was used. At each time point, a stack of 26 images separated by 0.5 μm was acquired, and the final time resolution is 16.8 sec/frame. Images were captured in 16-bit. Images were typically taken from the end of nc13 to the onset of gastrulation at nc14. During imaging, data acquisition was occasionally stopped for a few seconds to correct z-position, and data were concatenated afterwards. For each cross, three biological replicates were taken. The same laser power and microscope setting were used for each set of experiments.

### Plasmids

#### pbphi-DSCP_mTATA_-MS2-yellow-sna shadow enhancer

Two DNA oligos (5′-TTT CCC TCG AGG AGC TCG CCC GGG GAT CGA GCG CAG CGG GCG CCC CGG GCG CGG GGT GGC TGA GAG CAT CAG TTG TGA ATG AAT GTT CGA GCC GAG C-3′) and (5′-GGA AAG GAT CCG TTT GGT ATG CGT CTT GTG ATT CAA AGT TGG CTT ATT CAA AGG ATA TTA ACG AAG GCA GCG GCA CGT CTG CTC GGC TCG AAC AT-3′) were annealed and blunt-ended by PCR. Resulting DNA fragment was inserted between XhoI and BamHI sites of pbphi-DSCP-MS2-yellow (Yokoshi et al. 2020). Subsequently, a DNA fragment containing *sna* shadow enhancer was purified from pbphi-snail shadow enhancer (Lim et al. 2018) by digesting with HindIII and NheI. The resulting fragment was inserted between the HindIII and NheI sites of the plasmid.

#### pbphi-DSCP_mInr_-MS2-yellow-sna shadow enhancer

Two DNA oligos (5′-TTT CCC TCG AGG AGC TCG CCC GGG GAT CGA GCG CAG CGG TAT AAA AGG GCG CGG GGT GGC TGA GAG CAG TGA CAG TGA ATG AAT GTT CGA GCC GAG C-3′) and (5′-GGA AAG GAT CCG TTT GGT ATG CGT CTT GTG ATT CAA AGT TGG CTT ATT CAA AGG ATA TTA ACG AAG GCA GCG GCA CGT CTG CTC GGC TCG AAC AT-3′) were annealed and blunt-ended by PCR. Resulting DNA fragment was inserted between XhoI and BamHI sites of pbphi-DSCP-MS2-yellow (Yokoshi et al. 2020). Subsequently, a DNA fragment containing *sna* shadow enhancer was inserted between HindIII and NheI sites of the plasmid.

#### pbphi-DSCP_mMTE_-MS2-yellow-sna shadow enhancer

Two DNA oligos (5′-TTT CCC TCG AGG AGC TCG CCC GGG GAT CGA GCG CAG CGG TAT AAA AGG GCG CGG GGT GGC TGA GAG CAT CAG TTG TGA ATG AAT GTT ATC CAC GAG C-3′) and (5′-GGA AAG GAT CCG TTT GGT ATG CGT CTT GTG ATT CAA AGT TGG CTT ATT CAA AGG ATA TTA ACG AAG GCA GCG GCA CGT CTG CTC GTG GAT AAC AT-3′) were annealed and blunt-ended by PCR. Resulting DNA fragment was inserted between XhoI and BamHI sites of pbphi-DSCP-MS2-yellow (Yokoshi et al. 2020). Subsequently, a DNA fragment containing *sna* shadow enhancer was inserted between HindIII and NheI sites of the plasmid.

#### pbphi-DSCP_mDPE_-MS2-yellow-sna shadow enhancer

Two DNA oligos (5′-TTA AAC TCG AGG AGC TCG CCC GGG GAT CGA GCG CAG CGG TAT AAA AGG GCG CGG GGT GGC TGA GAG CAT CAG TTG TGA ATG AAT GTT CGA GCC GAG C-3′) and (5′-ACA TGG GAT CCG TTT GGT ATG CGT CTT GTG ATT CAA AGT TGG CTT ATT CAA AGG ATA TTA ACG AAG GCA GCG GCC ATG AGG CTC GGC TCG AAC ATT CAT T-3′) were annealed and blunt-ended by PCR. Resulting DNA fragment was inserted between XhoI and BamHI sites of pbphi-DSCP-MS2-yellow (Yokoshi et al. 2020). Subsequently, a DNA fragment containing *sna* shadow enhancer was inserted between HindIII and NheI sites of the plasmid.

#### pbphi-DSCP_mGAGA_-MS2-yellow-sna shadow enhancer

Two DNA oligos (5′-TTA AAC TCG AGG AGC TCG CCC GGG GAT CGA GCG CAG CGG TAT AAA AGG GCG CGG GGT GGC TCA CTG CAT CAG TTG TGA ATG AAT GTT CGA GCC GAG C-3′) and (5′-ACA TGG GAT CCG TTT GGT ATG CGT CTT GTG ATT CAA AGT TGG CTT ATT CAA AGG ATA TTA ACG AAG GCA GCG GCA CGT CTG CTC GGC TCG AAC ATT CAT T-3′) were annealed and blunt-ended by PCR. Resulting DNA fragment was inserted between XhoI and BamHI sites of pbphi-DSCP-MS2-yellow (Yokoshi et al. 2020). Subsequently, a DNA fragment containing *sna* shadow enhancer was inserted between HindIII and NheI sites of the plasmid.

#### pbphi-DSCP_3xZelda_-MS2-yellow-sna shadow enhancer

Two DNA oligos (5′-TTT CCC TCG AGC AGG TAG CCC GGG GAT CGA GCG CAG CGG TAT AAA AGG GCG CGG GGT GGC TGA GAG CAT CAG TTG TGA ATG AAT GTT CGA GCC GAG C-3′) and (5′-GGA AAG GAT CCG TTT GGT ATG CGT CTT GTG ATT CAA AGT TGG CTT ATT CAA AGG ATA TTA ACG AAG GCA GCG GCA CGT CTG CTC GGC TCG AAC AT-3′) were annealed and blunt-ended by PCR. Resulting DNA fragment was inserted between XhoI and BamHI sites of pbphi-DSCP-MS2-yellow (Yokoshi et al. 2020). Subsequently, two DNA oligos (5′-GGC CGC CAG GTA GCA GGT AGC-3′) and (5′-TCG AGC TAC CTG CTA CCT GGC-3′) were annealed and inserted into the plasmid using NotI and XhoI sites. Then, a DNA fragment containing *sna* shadow enhancer was inserted between HindIII and NheI sites of the plasmid.

#### lab_WT_-MS2-yellow-sna shadow enhancer

Two DNA oligos (5′-TTA AAG CGG CCG CGT CTG CAG AGG GGC GTG GCC AAG ACC AGC GGT TGT GCG GTC TGA AAG AAA CCG GGT TCG GGC CAG TAA TCA GTC AC-3′) and (5′-ACA TGG GAT CCC CGA AAA ACA CGA CTC CCG TTG GCG ATG ACG ACG ACG ACG TGC TGC CTG CGC GCT TAC CAA GTC GTG ACT GAT TAC TGG CCC GA-3′) were annealed and blunt-ended by PCR. Resulting DNA fragment was inserted between NotI and BamHI sites of pbphi-DSCP-MS2-yellow*-*sna shadow enhancer (Yokoshi et al. 2020).

#### lab_TATA_-MS2-yellow-sna shadow enhancer

Two DNA oligos (5′-TTA AAG CGG CCG CGT CTG CAG AGG GGC GTG GCC AAG ACC AGC GGT TGT GCG GTA TAA AAG AAA CCG GGT TCG GGC CAG TAA TCA GTC AC-3′) and (5′-ACA TGG GAT CCC CGA AAA ACA CGA CTC CCG TTG GCG ATG ACG ACG ACG ACG TGC TGC CTG CGC GCT TAC CAA GTC GTG ACT GAT TAC TGG CCC GA-3′) were annealed and blunt-ended by PCR. Resulting DNA fragment was inserted between NotI and BamHI sites of pbphi-DSCP-MS2-yellow*-*sna shadow enhancer (Yokoshi et al. 2020).

#### pbphi-DSCP_WT_-MS2-yellow-rhoNEE

A DNA fragment containing *rho*NEE was inserted between the HindIII and NheI sites of pbphi-DSCP_WT_-MS2-yellow-sna shadow enhancer. Sequence of *rho*NEE is the same as one used in the previous study (Fukaya et al. 2016).

#### pbphi-DSCP_mTATA_-MS2-yellow- rhoNEE

A DNA fragment containing *rho*NEE was inserted between the HindIII and NheI sites of pbphi-DSCP_mTATA_-MS2-yellow-sna shadow enhancer. Sequence of *rho*NEE is the same as one used in the previous study (Fukaya et al. 2016).

#### pbphi-DSCP_mInr_-MS2-yellow-rhoNEE

A DNA fragment containing *rho*NEE was inserted between the HindIII and NheI sites of pbphi-DSCP_mInr_-MS2-yellow-sna shadow enhancer. Sequence of *rho*NEE is the same as one used in the previous study (Fukaya et al. 2016).

#### pbphi-DSCP_mMTE_-MS2-yellow-rhoNEE

A DNA fragment containing *rho*NEE was inserted between the HindIII and NheI sites of pbphi-DSCP_mMTE_-MS2-yellow-sna shadow enhancer. Sequence of *rho*NEE is the same as one used in the previous study (Fukaya et al. 2016).

#### pbphi-DSCP_mDPE_-MS2-yellow-rhoNEE

A DNA fragment containing *rho*NEE was inserted between the HindIII and NheI sites of pbphi-DSCP_mDPE_-MS2-yellow-sna shadow enhancer. Sequence of *rho*NEE is the same as one used in the previous study (Fukaya et al. 2016).

#### pBS-attP-dsRed-SV40

Two DNA oligos (5 ́-TCG ACA GTT CTA GAC CCC CAA CTG AGA GAA CTC AAA GGT TAC CCC AGT TGG GGG AAT TCA TCG ATA ACT TCG TAT AAT GTA TGC TAT ACG AAG TTA TG-3 ́) and (5 ́-CTA GCA TAA CTT CGT ATA GCA TAC ATT ATA CGA AGT TAT CGA TGA ATT CCC CCA ACT GGG GTA ACC TTT GAG TTC TCT CAG TTG GGG GTC TAG AAC TG-3 ́) were annealed and inserted into the modified version of pBS-3xP3-dsRed plasmid using SalI and NheI sites.

#### pCFD3-dU6-ftz-1

Two DNA oligos (5′-GTC GCT GCA AGG ACA TTT CGC CGG-3′) and (5′-AAA CCC GGC GAA ATG TCC TTG CAG-3′) were annealed and inserted into the pCFD3-dU6:3gRNA vector (addgene # 49410) using BbsI sites.

#### pCFD3-dU6-ftz-2

Two DNA oligos (5′-GTC GCA ATT TGT GAA GAA GAG TCT-3′) and (5′-AAA CAG ACT CTT CTT CAC AAA TTG-3′) were annealed and inserted into the pCFD3-dU6:3gRNA vector (addgene # 49410) using BbsI sites.

#### pBS-ftz 5′Arm-attP-dsRed-SV40-ftz 3′Arm

A DNA fragment containing 3′ homology arm of *ftz* was amplified from genomic DNA using primers (5′-TTT AAA CTA GTT CTT GGG CAT GCT GCA ATT TG-3′) and (5′-TTA AAG CGG CCG CGA AGG GTA GGA TAG AAT CTG-3′), and digested with SpeI and NotI. The resulting fragment was inserted between the SpeI and NotI sites of pBS-attP-dsRed-SV40. Subsequently, a DNA fragment containing 5′ homology arm of *ftz* was amplified from genomic DNA using primers (5′-TTT AAG GTA CCA TGA AGA TCC TAC GCT GTG C-3′) and (5′-TTA AAG TCG ACG CGA AAT GTC CTT GCA GGC ACG-3′), and digested with KpnI and SalI. The resulting fragment was inserted between the KpnI and SalI sites of the plasmid.

#### pbphi-ftz WT core promoter-ftz-HA-24xMS2-aTub 3′UTR

A DNA fragment containing *ftz* transcription unit fused with C-terminal HA-tag was amplified from genomic DNA using primers (5′-TTT AAG GAT CCA CTA GTA TGG CCA CCA CAA ACA GCC AGA GC-3′) and (5′-TTA AAG GAT CCT CAT GCA TAA TCC GGA ACA TCA TAC GGA TAA GAC AGA TGG TAG AGG TCC TGT GG-3′), and digested with BamHI. The resulting fragment was inserted between the BglII and BamHI sites of the pbphi-hbP2 promoter-lacZ-24xMS2-αTub 3 ́UTR (modified version of pbphi-hbP2 promoter-lacZ-24xPP7-αTub 3 ́UTR described in (Fukaya et al. 2017)). Subsequently, a DNA fragment containing native *ftz* core promoter sequence was amplified from genomic DNA using primers (5′-TTT AAA AGC TTT GTC ATG CGC AGG GAT ATT TAT GCG-3′) and (5′-TTA AAA CTA GTA TCG GAT GTG TAT TGC TAG ATT TC-3′), and digested with HindIII and SpeI. The resulting fragment was inserted between the HindIII and SpeI sites of the plasmid.

#### pbphi-ftz mTATA core promoter-ftz-HA-24xMS2-aTub 3′UTR

Two DNA oligos (5′-ACT ATA AGC TTT GTC ATG CGC AGG GAT ATT TAT GCG CTA TAA CGC CGA GCG TGT GCC GAG GGC TCT CTG ATT TTG CGC GCC CCG CAG GAT CTG CCG CAG G-3′) and (5′-AAT CCA CTA GTA TCG GAT GTG TAT TGC TAG ATT TCT TCT CTA ACT CTG CGA TGT GCA CGC AAC GCT GGT GAG TTT GCG AAT GAG CTG GTC CTG CGG CAG ATC CTG CGG G-3′) were annealed and blunt-ended by PCR. The resulting fragment was inserted between the HindIII and SpeI sites of pbphi-ftz WT core promoter-ftz-HA-24xMS2-aTub 3′UTR.

#### pbphi-ftz mDPE core promoter-ftz-HA-24xMS2-aTub 3′UTR

Two DNA oligos (5′-ACA TGA AGC TTT GTC ATG CGC AGG GAT ATT TAT GCG CTA TAA CGC CGA GCG TGT GCC GAG GGC TCT CTG ATT TTG CTA TAT ATG CAG GAT CTG CCG CAG G-3′) and (5′-AAT CCA CTA GTA TCG GAT GTG TAT TGC TAG ATT TCT TCT CTA ACT CTG CAT GAG GCA CGC AAC GCT GGT GAG TTT GCG AAT GAG CTG GTC CTG CGG CAG ATC CTG CAT A-3′) were annealed and blunt-ended by PCR. The resulting fragment was inserted between the HindIII and SpeI sites of pbphi-ftz WT core promoter-ftz-HA-24xMS2-aTub 3′UTR.

### Image analysis

All the image processing methods and analysis were implemented in MATLAB (R2019b, MathWorks).

### Segmentation of nuclei

For each time point, maximum projections were obtained for all 26 z-sections per image. His2Av-mRFP was used to segment nuclei. 512 × 512 maximum projection images were initially cropped into 430 × 430 (Figure 1, 4, 6) or 300 × 430 (Figure 2 and 3) to remove nuclei at the edge, and used for subsequent analysis. In the analysis of *rho*NEE reporters (Figure S2), 512 × 512 maximum projection images were initially cropped into 300 × 500 to remove nuclei outside of the expression domain. For nuclei segmentation, we used two different methods. In the first method, His2Av images were pre-processed with Gaussian filtering, top-hap filtering, and adaptive histogram equalization, in order to enhance the signal-to-noise contrast. Processed images were converted into binary images using a threshold value obtained from Otsu’s method. Nuclei were then watershedded to further separate and distinguish from neighboring nuclei. Subsequently, binary images were manually corrected using Fiji (https://fiji.sc). In the second method, His2Av images were first blurred with Gaussian filter to generate smooth images. Pixels expressing intensity higher than 5% of the global maxima in the histogram of His2Av channel were removed. Processed images were converted into binary images using a custom threshold-adaptative segmentation algorithm. Threshold values were determined at each time frame by taking account of 1) histogram distribution of His2Av channel and 2) the number and size of resulting connected components. Boundaries of components were then modified to locate MS2 transcription dots inside of nearest nuclei. In brief, pixels with intensity twice larger than mean intensity of MS2 channel were considered as transcription dots, and new binary images were created for each time frame. The Euclidean distances between the centroid of binarized transcription dot and all boundaries of segmented nuclei were calculated. Boundary of the nucleus with the smallest Euclidean distance was modified in order to capture transcription dot within a nucleus. Centroids of connected components in nuclei segmentation channel were used to compute the Voronoi cells of the image. Resulting binary images were manually corrected by using Fiji (https://fiji.sc).

### Tracking of nuclei

Nuclei tracking was done by finding the object with minimal movement across the frames of interest. For each nucleus in a given frame, the Euclidean distances between the centroids of the nucleus in current time frame and the nuclei in the next or previous time frame were determined. The nucleus with the minimum Euclidean distance was considered as the same lineage.

### Recording of MS2 signals

Maximum projections of raw images were used to record MS2 fluorescence intensities. Using segmented regions, fluorescence intensities within each nucleus were extracted. Signals of MS2 transcription dots were determined by taking an average of the top three pixels with the highest fluorescence intensity within each nucleus. After obtaining MS2 trajectories, median fluorescence intensities within a nucleus were subtracted as a background. In Figure S2, signals of MS2 transcription dots were determined by taking integral in a 5 × 5 pixel region centering the brightest pixel within a nucleus after subtracting median fluorescence intensity within a nucleus as a background. Subsequently, minimum MS2 intensities were determined for individual trajectories and subtracted to make the baseline zero.

### Detection of transcriptional bursting

A transcriptional burst was defined as a local change in fluorescence intensity. First, signal trajectories were smoothed by the moving average method. When a nucleus had above-threshold transcription activity, burst was considered to be started. Burst was considered to be ended when the intensity dropped below 55% of the local peak value. When the burst duration is less than 5 timeframes, it was considered as a false-positive derived from detection noise. When signal trace exhibited continuous decreasing at the beginning of burst detection, it was also not considered as a burst. Location of defined burst was then moved two timeframes afterwards to better capture the center of individual bursting event. Same method and threshold value were used for each set of experiments.

### Description of bursting properties

From each trajectory, number of bursts, amplitude and duration of each burst, and total integrated signal (output) produced by each nucleus were measured. To determine amplitude, the peak value during the burst was measured using trajectories after smoothing. The duration was determined by measuring the length of each burst. Total RNA production was measured by taking the area under the raw trajectory. The amplitude and duration for each nucleus were determined by taking average of all analyzed bursts in a single nucleus. The probability of burst induction was determined as a frequency divided by a length of time after the initial burst for each active nucleus.

### Fraction of instantaneously and cumulative active nuclei

For each time frame, nuclei with MS2 intensity above the threshold were considered as active. Threshold was determined for each group by calculating the 10% of the maximum MS2 intensity across all trajectories within a group.

### Mean MS2 intensity per actively transcribing nucleus

MS2 intensities in instantaneously active nuclei at each time point were averaged.

### Computational reconstitution of *ftz* expression

Newly synthesized *ftz* mRNAs were considered to be linearly degraded with a half-life of 7 min according to previous measurements in early embryos (Edgar et al. 1986). Amount of *ftz* mRNA remained to be undegraded at the end of analysis was estimated using live imaging data. Using segmentation mask, individual nuclei were false-colored with the pixel intensity proportional to the level of intact *ftz* mRNA in a given nucleus. Resulting image was then colored and layered over the maximum projected image of His2Av-mRFP.

### False-coloring by instantaneous MS2 signals

MS2 signal intensity at each nucleus was measured as described in the previous section. Using segmentation mask, individual nuclei were false-colored with the pixel intensity proportional to the instantaneous MS2 signal at given time in a given nucleus. Resulting image was then colored and layered over the maximum projected image of His2Av-mRFP.

