## Supplemental Figure for "Dynamic modulation of enhancer responsiveness by core promoter elements in living *Drosophila* embryo"

Yokoshi_Figure S1

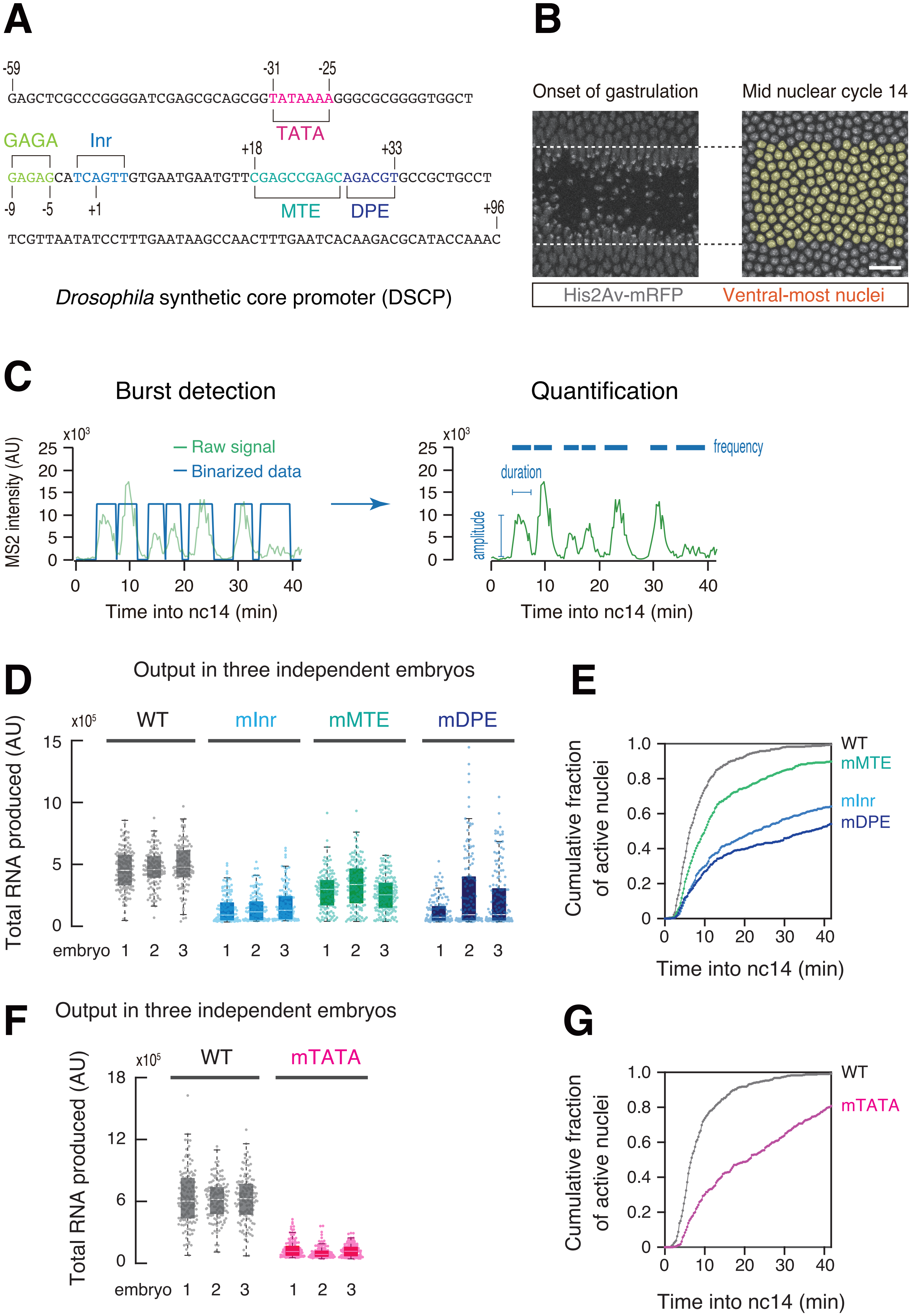

**Figure S1. Quantification of transcriptional bursting.**

(A) Overall structure of the *Drosophila* synthetic core promoter (DSCP).

(B) Snapshot of a gastrulating embryo (left) and false-coloring of ventral-most nuclei at mid nc14 (right). The maximum projected image of a histone marker (His2Av-mRFP) is shown in gray. Images are oriented with ventral view facing up. Scale bar indicates 20 μm.

(C) Trajectory of transcription activity shown in Figure 1D (left). Amplitude and duration were measured for every single bursting event. Frequency was determined by counting total number of bursts during the analysis.

(D) Boxplots showing the distribution of total output in each of three independent embryos. The box indicates the lower (25%) and upper (75%) quantile and the solid line indicates the median. Whiskers extend to the most extreme, non-outlier data points. A number of nuclei analyzed for the reporter genes is as follows. WT; 147, 129 and 127. mInr; 150, 145 and 140. mMTE; 156, 158 and 195. mDPE; 145, 135 and 164. AU; arbitrary unit.

(E) Cumulative fraction of actively transcribing nuclei at the ventral region. A total of 403, 435, 509 and 444 ventral-most nuclei were analyzed from three individual embryos for WT, mInr, mMTE and mDPE DSCP.

(F) Boxplots showing the distribution of total output in each of three independent embryos. The box indicates the lower (25%) and upper (75%) quantile and the solid line indicates the median. Whiskers extend to the most extreme, non-outlier data points. A number of nuclei analyzed for the reporter genes is as follows. WT; 133, 140 and 128. mTATA; 139, 139 and 128. AU; arbitrary unit.

(G) Cumulative fraction of actively transcribing nuclei at the ventral region. A total of 401 and 406 ventral-most nuclei were analyzed from three individual embryos for WT and mTATA DSCP.

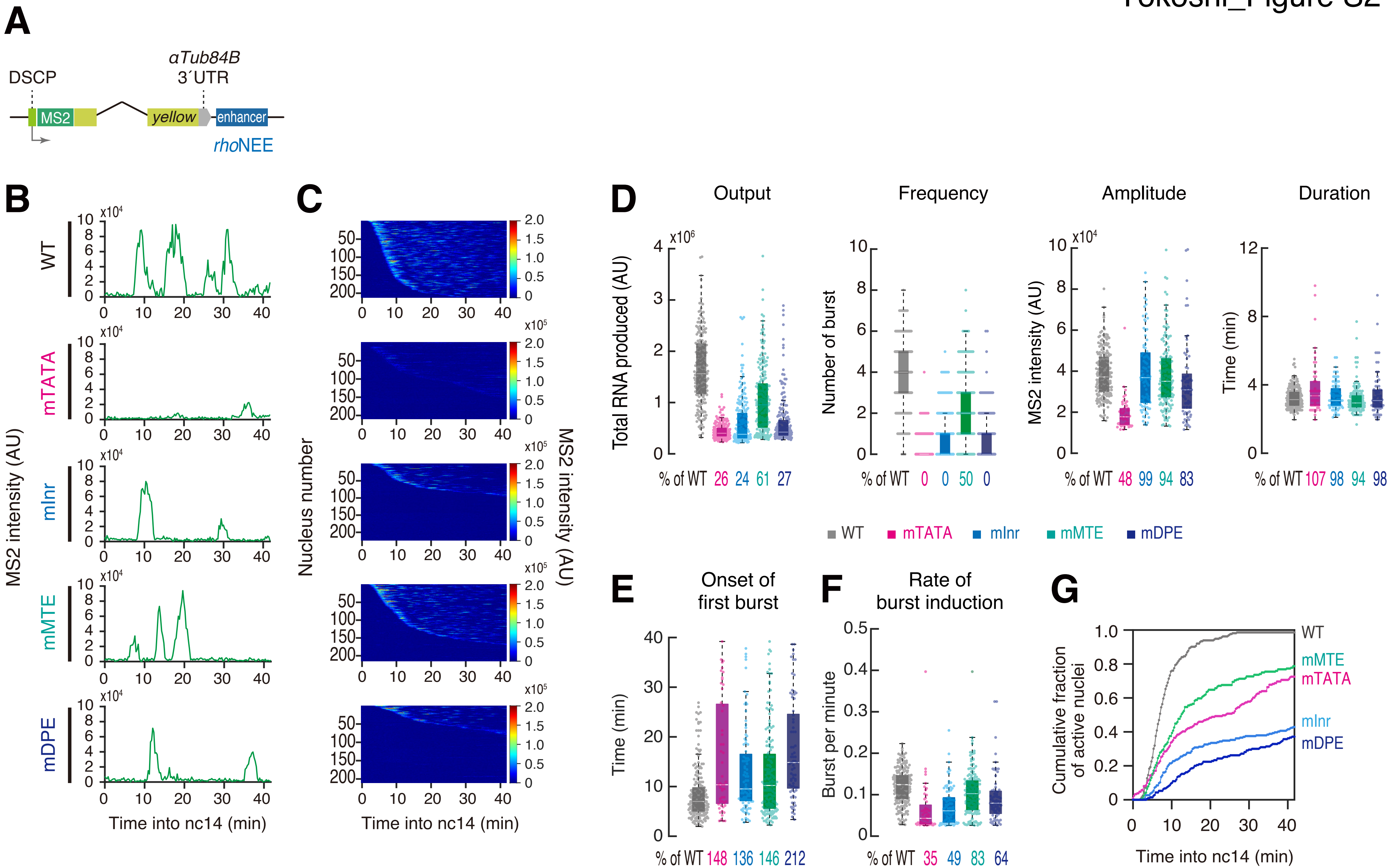
A
αTub84B
3´UTR
DSCP
MS2
yellow
enhancer
rhoNEE
B
WT
mTATA
mInr
mMTE
mDPE
MS2 intensity (AU)
Time into nc14 (min)
C
Nucleus number
MS2 intensity (AU)
Time into nc14 (min)
D
Output
Frequency
Amplitude
Duration
Total RNA produced (AU)
Number of burst
MS2 intensity (AU)
Time (min)
% of WT 26 24 61 27
% of WT 0 0 50 0
% of WT 48 99 94 83
% of WT 107 98 94 98
WT mTATA mInr mMTE mDPE
E
Onset of first burst
Time (min)
% of WT 148 136 146 212
F
Rate of burst induction
Burst per minute
% of WT 35 49 83 64
G
Cumulative fraction of active nuclei
Time into nc14 (min)
WT
mMTE
mTATA
mInr
mDPE

**Figure S2. Live imaging analysis of reporter genes under the control of *rho*NEE.**

(A) Schematic representation of the *yellow* reporter gene containing the 155-bp DSCP, the 666-bp *rho*NEE, and 24x MS2 RNA stem loops within the 5´ UTR.

(B) A representative trajectory of transcription activity of the MS2 reporter genes with WT (top), mTATA (upper middle), mInr (middle), mMTE (lower middle) and mDPE DSCP (bottom) in individual nuclei. AU; arbitrary unit.

(C) MS2 trajectories for all analyzed nuclei. Each row represents the MS2 trajectory for a single nucleus. A total of 213, 211, 224, 216 and 210 nuclei located at the expression domain, respectively, were analyzed from three independent embryos for the reporter genes with WT (top), mTATA (upper middle), mInr (middle), mMTE (lower middle) and mDPE DSCP (bottom). Nuclei were ordered by their onset of transcription in nc14. AU; arbitrary unit.

(D) Boxplots showing the distribution of total output (left), burst frequency (middle left), burst amplitude (middle right) and burst duration (right). The box indicates the lower (25%) and upper (75%) quantile and the solid line indicates the median. Whiskers extend to the most extreme, non-outlier data points. A total of 213, 211, 224, 216 and 210 nuclei located at the expression domain, respectively, were analyzed from three independent embryos for the reporter genes with WT, mTATA, mInr, mMTE and mDPE DSCP. Median values relative to the WT reporter are shown at the bottom. AU; arbitrary unit.

(E and F) Boxplots showing the distribution of the timing of first burst (E) and burst frequency normalized by the length of time after the first burst (F). The box indicates the lower (25%) and upper (75%) quantile and the solid line indicates the median. Whiskers extend to the most extreme, non-outlier data points. A total of 213, 211, 224, 216 and 210 nuclei, respectively, were analyzed from three independent embryos for the reporter genes with WT, mTATA, mInr, mMTE and mDPE DSCP. Median values relative to the WT reporter are shown at the bottom.

(G) Cumulative fraction of actively transcribing nuclei at the ventral neurogenic ectoderm. A total of 213, 211, 224, 216 and 210 nuclei, respectively, were analyzed from three independent embryos for the reporter genes with WT, mTATA, mInr, mMTE and mDPE DSCP.

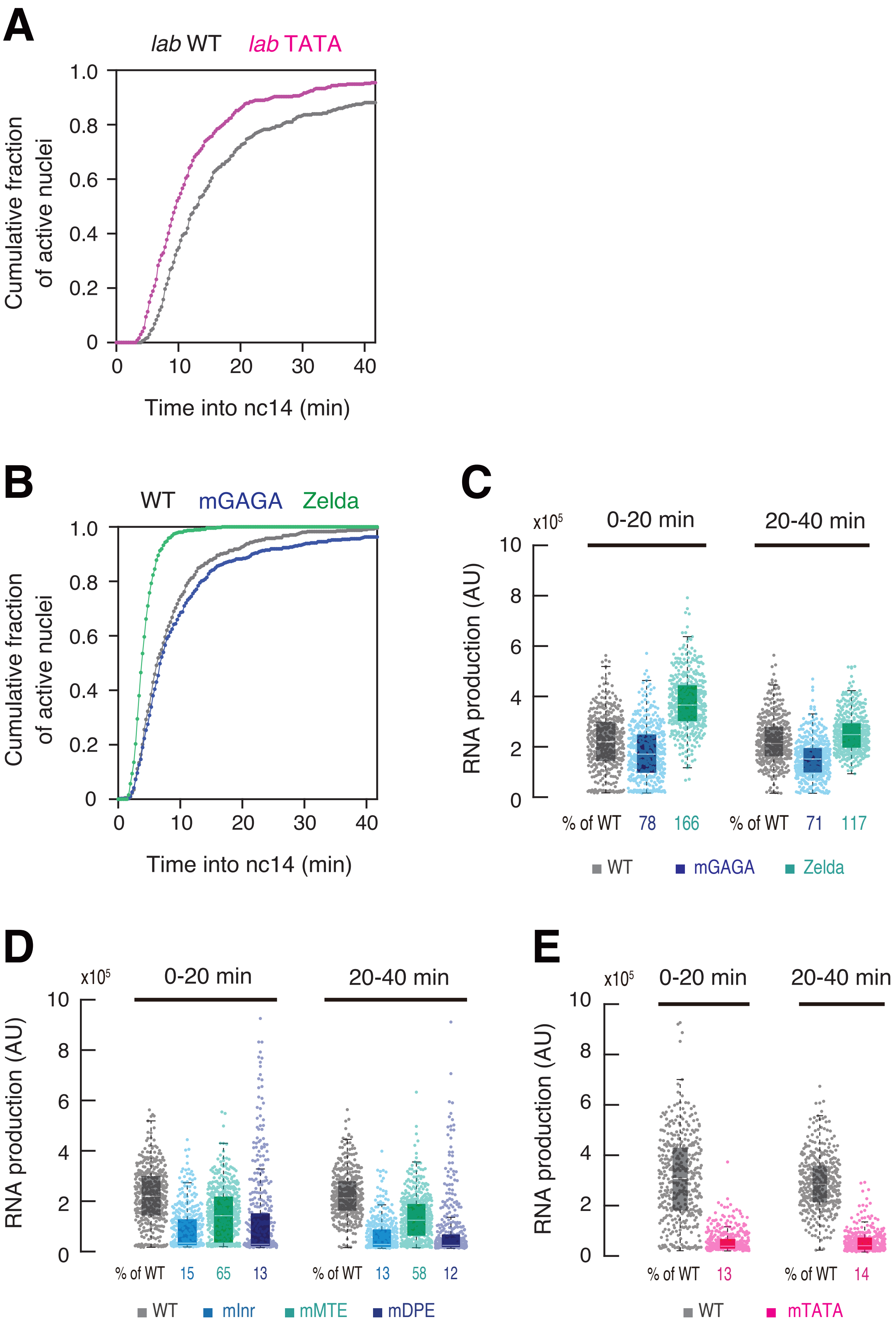
A
lab WT
lab TATA
Cumulative fraction of active nuclei
1.0
0.8
0.6
0.4
0.2
0
0
10
20
30
40
Time into nc14 (min)
B
WT
mGAGA
Zelda
Cumulative fraction of active nuclei
Time into nc14 (min)
C
x10^5
0-20 min
20-40 min
RNA production (AU)
% of WT 78 166
% of WT 71 117
WT
mGAGA
Zelda
D
x10^5
0-20 min
20-40 min
RNA production (AU)
% of WT 15 65 13
% of WT 13 58 12
WT
mInr
mMTE
mDPE
E
x10^5
0-20 min
20-40 min
RNA production (AU)
% of WT 13
% of WT 14
WT
mTATA

**Figure S3. Zelda facilitates induction of the first round of transcription.**

(A) Cumulative fraction of actively transcribing nuclei at the ventral region. A total of 336 and 371 ventral-most nuclei were analyzed from three independent embryos for the reporter genes with unmodified TATA-less and modified TATA-containing *lab* core promoter.

(B) Cumulative fraction of actively transcribing nuclei at the ventral region. A total of 403, 458 and 458 ventral-most nuclei were analyzed from three individual embryos for WT, mGAGA and Zelda DSCP. Plot of WT is the same as the plot in Figure S1E.

(C) Boxplots showing the distribution of output during first 0-20 min (left) and last 20-40 min of the analysis (right). The box indicates the lower (25%) and upper (75%) quantile and the solid line indicates the median. Whiskers extend to the most extreme, non-outlier data points. A total of 403, 458 and 458 ventral-most nuclei, respectively, were analyzed from three independent embryos for the reporter genes with WT, mGAGA and Zelda DSCP. Median values relative to the WT reporter are shown at the bottom. AU; arbitrary unit.

(D) Boxplots showing the distribution of output during first 0-20 min (left) and last 20-40 min of the analysis (right). The box indicates the lower (25%) and upper (75%) quantile and the solid line indicates the median. Whiskers extend to the most extreme, non-outlier data points. A total of 403, 435, 509 and 444 ventral-most nuclei, respectively, were analyzed from three independent embryos for the reporter genes with WT, mInr, mMTE and mDPE DSCP. Median values relative to the WT reporter are shown at the bottom. Plot of WT is the same as the plot in Figure S3C. AU; arbitrary unit.

(E) Boxplots showing the distribution of output during first 0-20 min (left) and last 20-40 min of the analysis (right). The box indicates the lower (25%) and upper (75%) quantile and the solid line indicates the median. Whiskers extend to the most extreme, non-outlier data points. A total of 401 and 406 ventral-most nuclei, respectively, were analyzed from three independent embryos for the reporter genes with WT and mTATA DSCP. Median values relative to the WT reporter are shown at the bottom. AU; arbitrary unit.

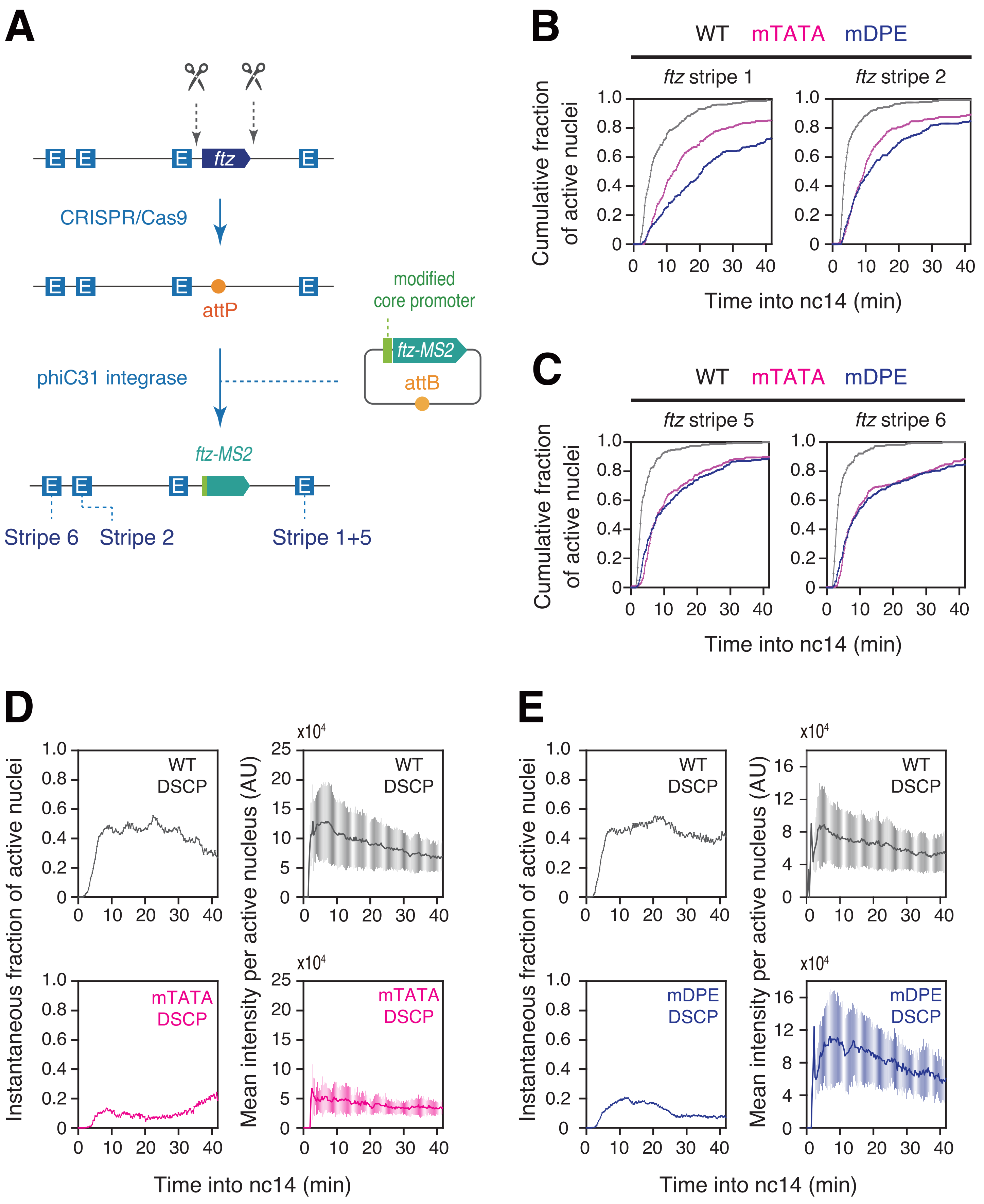
A
ftz
CRISPR/Cas9
attP
modified
core promoter
ftz-MS2
attB
phiC31 integrase
ftz-MS2
Stripe 6
Stripe 2
Stripe 1+5
B
WT mTATA mDPE
ftz stripe 1
ftz stripe 2
Cumulative fraction of active nuclei
Time into nc14 (min)
C
WT mTATA mDPE
ftz stripe 5
ftz stripe 6
Cumulative fraction of active nuclei
Time into nc14 (min)
D
WT DSCP
mTATA DSCP
Instantaneous fraction of active nuclei
Mean intensity per active nucleus (AU)
x10[4]
Time into nc14 (min)
E
WT DSCP
mDPE DSCP
Instantaneous fraction of active nuclei
Mean intensity per active nucleus (AU)
x10[4]
Time into nc14 (min)

**Figure S4. Dynamics of *ftz* transcription at the engendered alleles.**

(A) Schematic representation of the core promoter modification approach. Approximate locations of enhancers regulating *ftz* expression in early embryos are indicated as "E". Note that extra sequences derived from plasmid backbone, ampicillin resistant gene (*ampR*) and mini-white marker gene (*miniW*) were incorporated into the *ftz* locus after phiC31-mediated integration.

(B) Cumulative fraction of actively transcribing nuclei at anterior stripe 1/2. A total of 234, 236 and 228 nuclei at stripe 1, and 248, 241 and 242 nuclei at stripe 2 were analyzed from three individual embryos for the *ftz-MS2* with WT, mTATA and mDPE core promoter, respectively.

(C) Cumulative fraction of actively transcribing nuclei at posterior stripe 5/6. A total of 243,238 and 240 nuclei at stripe 5, and 232, 243 and 240 nuclei at stripe 6 were analyzed from three individual embryos for the *ftz-MS2* with WT, mTATA and mDPE core promoter, respectively.

(D) Instantaneous fraction of actively transcribing nuclei (left) and mean MS2 intensity per actively transcribing nucleus (right). A total of 401 and 406 ventral-most nuclei, respectively, were analyzed from three independent embryos for the reporter genes with WT and mTATA DSCP. AU; arbitrary unit.

(E) Instantaneous fraction of actively transcribing nuclei (left) and mean MS2 intensity per actively transcribing nucleus (right). A total of 403 and 444 ventral-most nuclei, respectively, were analyzed from three independent embryos for the reporter genes with WT and mDPE DSCP. AU; arbitrary unit.

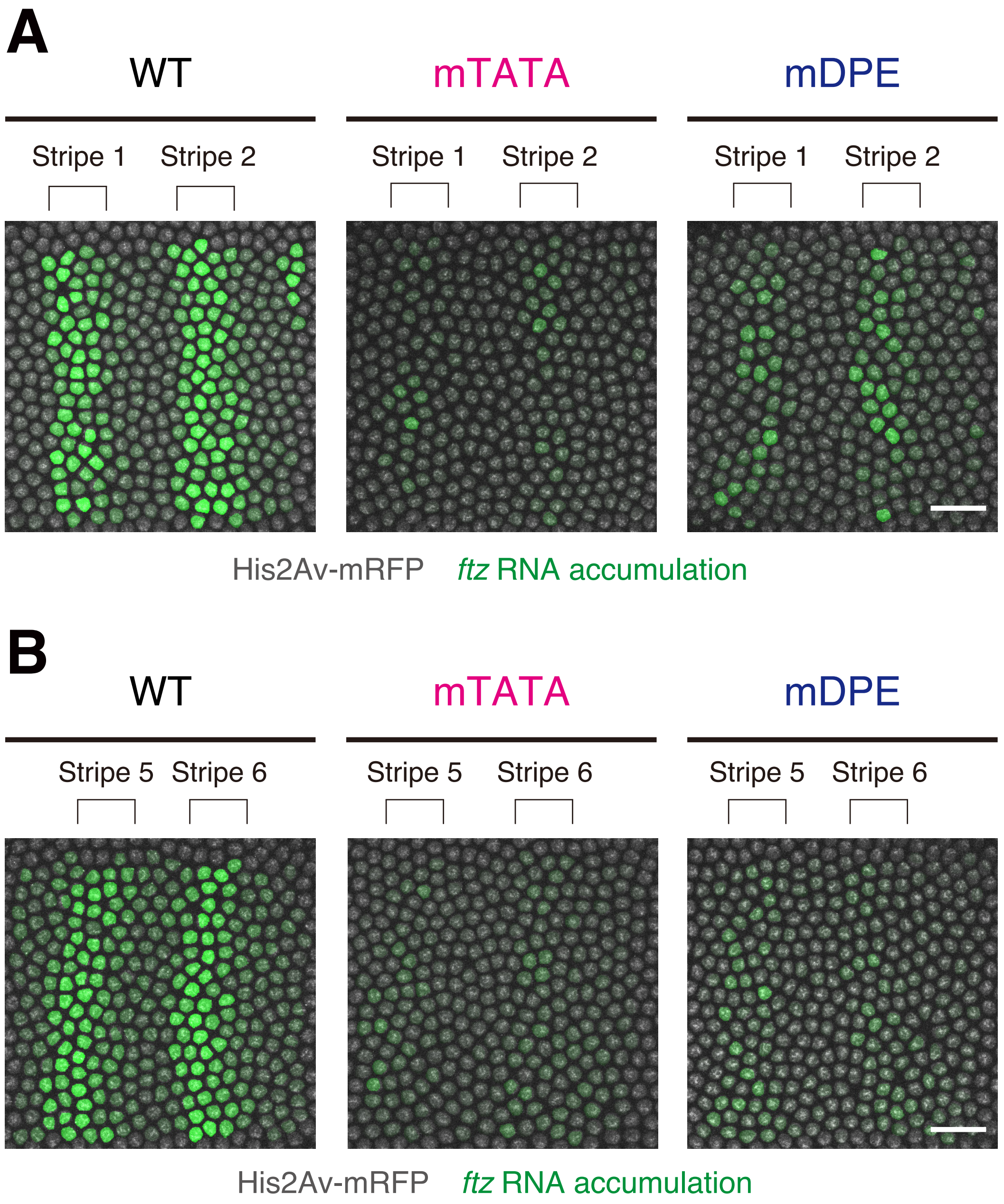
A
WT
mTATA
mDPE
Stripe 1
Stripe 2
Stripe 1
Stripe 2
Stripe 1
Stripe 2
His2Av-mRFP
*ftz* RNA accumulation
B
WT
mTATA
mDPE
Stripe 5
Stripe 6
Stripe 5
Stripe 6
Stripe 5
Stripe 6
His2Av-mRFP
*ftz* RNA accumulation

**Figure S5. TATA and DPE mutations lead to sporadic stripe formation.**

(A) Computational reconstitution of *ftz* mRNA accumulation at the anterior stripe region. Scale bar indicates 20 μm.

(B) Computational reconstitution of *ftz* mRNA accumulation at the posterior stripe region. Scale bar indicates 20 μm.

### Movie Legends

**Movie S1. Live imaging of the MS2 reporters with WT, mInr, mMTE and mDPE DSCP.**

Live imaging of *MS2-yellow-sna shadow enhancer* containing WT (left), mInr (left middle), mMTE (right middle) and mDPE DSCP (right) during nc14. Each nucleus was false-colored with the intensity proportional to the instantaneous MS2 signal at given time in a given nucleus. The maximum projected image of His2Av-mRFP is shown in gray. Images are oriented with ventral view facing up.

**Movie S2. Live imaging of the MS2 reporters with WT and mTATA DSCP.**

Live imaging of *MS2-yellow-sna shadow enhancer* containing WT (left) and mTATA DSCP (right) during nc14. Each nucleus was false-colored with the intensity proportional to the instantaneous MS2 signal at given time in a given nucleus. The maximum projected image of His2Av-mRFP is shown in gray. Images are oriented with ventral view facing up.

**Movie S3. Live imaging of the MS2 reporter with WT and TATA *lab* core promoter.**

Live imaging of *MS2-yellow-sna shadow enhancer* containing unmodified TATA-less (left) and modified TATA-containing *lab* core promoter (right) during nc14. Each nucleus was false-colored with the intensity proportional to the instantaneous MS2 signal at given time in a given nucleus. The maximum projected image of His2Av-mRFP is shown in gray. Images are oriented with ventral view facing up.

**Movie S4. Live imaging of the MS2 reporters with WT, mGAGA and Zelda DSCP.**

Live imaging of *MS2-yellow-sna shadow enhancer* containing WT (left), mGAGA (middle) and Zelda DSCP (right) during nc14. Each nucleus was false-colored with the intensity proportional to the instantaneous MS2 signal at given time in a given nucleus. The maximum projected image of His2Av-mRFP is shown in gray. Images are oriented with ventral view facing up.

**Movie S5. Live imaging of *ftz-MS2* complementation alleles.**

Live imaging of *ftz-MS2* containing WT (top), mTATA (middle) and mDPE core promoter (bottom) during nc14. Anterior stripe 1 and 2 regions were shown in left. Posterior stripe 5 and 6 regions were shown in right. Each nucleus was false-colored with the intensity proportional to the instantaneous MS2 signal at given time in a given nucleus. The maximum projected image of His2Av-mRFP is shown in gray. Images are oriented with anterior to the left.
